# Curcumin and Sulforaphane Preserve Mobility in Aging *Caenorhabditis elegans* via Distinct yet Complementary Transcriptional Signatures

**DOI:** 10.64898/2026.06.23.734065

**Authors:** R. P. Vivek-Ananth, Durai Sellegounder, Karthik Mohanraj, Sushmita Maitra, Chris Saunter, David Weinkove, Eric Verdin, Stephen M. Phipps, Nathan D. Price

## Abstract

Aging involves a progressive decline in bodily functions, underscoring the need for interventions that enhance healthspan. In this study, we screened nine natural products in *Caenorhabditis elegans* using whole-organism phenotyping to assess mobility endpoints, and subsequently focused on curcumin, sulforaphane, and their combination. In replicated follow-up experiments, all three interventions improved late-adult mobility after Day 2 of adulthood. Sulforaphane and the combination provided the strongest gains, whereas curcumin showed a distinct benefit profile, with more pronounced effects on time active measures than on speed-based metrics. To examine associated molecular changes, we performed transcriptomic profiling on Day 3 adults. Curcumin was associated with lipid and sphingolipid remodeling together with reduced expression of several innate immune effectors, whereas sulforaphane induced glutathione-linked detoxification signatures involving multiple *gst* genes. The combination retained major features of both single-compound responses while adding combination-specific changes that broadened detoxification-associated signatures and extended repression of lectin-and lysozyme-associated genes. Transcription factor activity inference further supported SKN-1-linked detoxification responses under sulforaphane and the combination. Overall, these results suggest that curcumin and sulforaphane engage distinct yet partially convergent maintenance-related programs, and that their combination broadens the underlying molecular response without producing additive mobility gains. These findings motivate further testing of natural product combinations in healthspan-related contexts.

## Introduction

Aging represents a complex biological phenomenon marked by a gradual decline in physiological functions, increasing susceptibility to age-related diseases (World Health Organization 2015; López-Otín et al. 2013; Niccoli and Partridge 2012). This universal phenomenon occurs across nearly all multicellular organisms, with biological functions deteriorating in various organs during the post-reproductive phase (Jones et al. 2014; Kirkwood and Austad 2000). The global aging population is growing rapidly, with individuals aged 60 and older projected to double their current proportion by 2050 (World Health Organization 2015). This demographic shift poses significant challenges for healthcare systems and highlights the need for interventions that not only extend lifespan but also enhance healthspan, the period of life spent in good health (Kennedy et al. 2014).

Natural products and dietary supplements offer a promising avenue for enhancing healthspan (Argyropoulou et al. 2013; Bjørklund et al. 2022). Several natural products derived from plants and plant extracts have a long history of use in traditional medicine for treating ailments associated with aging (Argyropoulou et al. 2013; Mohanraj et al. 2018; Vivek-Ananth et al. 2023). Scientific investigations have shown that specific compounds within the natural products possess antioxidant and anti-inflammatory properties that may combat age-related health decline (Wink 2022; Lin et al. 2023; Davinelli et al. 2025). For instance, compounds like curcumin (from turmeric), sulforaphane (from cruciferous vegetables), resveratrol (from grapes and berries), quercetin (from various fruits and vegetables), and catechins (from green tea) have been shown to modulate multiple pathways associated with aging (Lin et al. 2023; Davinelli et al. 2025; Shen et al. 2013; J. Xu et al. 2023; Ji et al. 2021). These include activating the Nrf2/SKN-1 pathway, a key regulator of cellular defense mechanisms, and reducing oxidative stress and inflammation, which are major contributors to age-related damage (Argyropoulou et al. 2013; Lin et al. 2023; Ogawa et al. 2016; Fang et al. 2017; Blackwell et al. 2015).

The nematode *Caenorhabditis elegans* has become a pivotal model organism for aging research, particularly for evaluating anti-aging interventions such as natural products (Weinkove and Zavagno 2021; Kenyon 2010; Kirchweger et al. 2023). Its unique experimental advantages, including a short lifespan, ease of cultivation, and genetic tractability enable rapid and cost-effective screening of potentially beneficial compounds (Lin et al. 2023; Kaletta and Hengartner 2006; O’Reilly et al. 2014). Further, the mechanisms of aging, and key longevity pathways including insulin signaling, proteostasis, and oxidative stress responses, are highly conserved between *C. elegans* and humans, making it a valuable model for studying the molecular basis of aging (Lin et al. 2023; Blackwell et al. 2015; Kenyon 2010). Recent advancements in automated phenotyping have enhanced its utility in screening studies, allowing for the real-time, non-invasive monitoring of healthspan markers across large populations (Weinkove and Zavagno 2021; Zavagno et al. 2024; Mathew et al. 2012; Churgin et al. 2017). Moreover, *C. elegans* is particularly suited for studying the multi-target effects of natural products, as it integrates systemic responses to complex interventions (Kirchweger et al. 2023; Wink 2022).

This study initially evaluated nine selected natural products, plant extracts, and dietary supplements (berberine, bergamot extract, curcumin, ginkgo, liquorice root extract, *Rhodiola rosea* root extract, *Salvia miltiorrhiza* root extract, and sulforaphane; nutritional supplement nicotinamide riboside) for their potential to enhance healthspan in *C. elegans*, using automated whole-organism phenotyping. We tracked five mobility endpoints: fraction of moving worms (fraction moving), speed of moving worms, and speed of all worms (population-averaged speed), hours moved, and distance moved. From this screen, we prioritized curcumin and sulforaphane for deeper investigation of their effects on the healthspan of *C. elegans*. To the best of our knowledge, this represents the first such report to study the combined effect of curcumin and sulforaphane on *C. elegans* healthspan. Furthermore, we integrated these healthspan phenotypes with transcriptomic profiles to gain novel insights into the aging-related molecular pathways modulated by curcumin and sulforaphane, individually and in combination.

## Methods

### *C. elegans* Culture and Maintenance

Worms were obtained from the *Caenorhabditis elegans* Genetics Center (University of Minnesota, Minneapolis, MN). Wild-type (WT) N2 strain was used for development and reproductive toxicity assays, SS104 *glp-4*(bn2) strain was used for the healthspan assay. SS104 *glp-4*(bn2) strain was maintained at 15°C and moved to 24°C at L2/L3 stage to induce sterility. All worms were maintained in Defined Medium (DM) plates seeded with *Escherichia coli* OP50 as food source. Care was taken to keep worms in a fed condition for several generations to avoid artifacts due to starvation. The bacterial lawns with the overlaid compounds were allowed to partially dry at room temperature and then transferred to a 24°C incubator to dry overnight.

### Natural Compound Solution and Plate Preparation

The natural products, plant extracts, and supplements used in this study were purchased from several vendors. Curcumin phytosome, a formulation of curcumin, ginkgo phytosome, a formulation made from the *Ginkgo biloba* leaf extract, berberine phospholipid formulation, liquorice root extract, and *Rhodiola rosea* root extract were purchased from Indena. Sulforaphane, *Salvia miltiorrhiza* root extract, bergamot extract from *Citrus bergamia*, and Nicotinamide Riboside Hydrogen L-Malate were purchased from MedKoo, Nura, Bernett and Biosynth-Carbosynth respectively. Sulforaphane was stored at 4° C, and the rest of the compounds were stored at room temperature in powder form.

Stock suspensions or solutions (50 mg/mL) of the natural compounds were prepared in sterile water. The suspension was left at room temperature for 1–2 hours and vortexed gently every 30 mins for the compounds to dissolve completely and diluted to the desired concentration. The suspensions were briefly centrifuged at 2000 rpm for 30 seconds to remove large insoluble debris from the natural products but allowing small particles < 1 µm that could be ingested by *C. elegans* to remain. 80µL of the supernatant was carefully added to the lawn and allowed to dry overnight at incubator on the bacterial lawn seeded on DM assay plates.

### Qualitative Developmental and Reproductive Screen

We performed a qualitative pre-screen to identify obvious developmental or reproductive impairment at the concentrations used for the initial compound screen. Four WT N2 L4-stage worms were placed on each DM plate containing the indicated intervention and maintained at 24°C. Progeny production and growth were assessed qualitatively on Days 3 and 5, time points at which second-generation animals become apparent under normal development. Bacterial lawn consumption was also monitored qualitatively as an additional gross readout of worm growth and plate health. This screen was intended to identify obvious developmental or reproductive defects rather than provide a quantitative assessment of fertility or developmental rate. Bacterial lawns were visually comparable to control for all interventions except sulforaphane. Because sulforaphane at 10 mg/mL produced a visibly thinner OP50 lawn, we performed an OD600-based bacterial growth assay across lower sulforaphane concentrations; growth inhibition was detected at 2.5 and 10 mg/mL but not at concentrations ≤ 0.625 mg/mL, and 0.625 mg/mL was therefore used in follow-up experiments.

### Plate Preparation for Healthspan Assay

On Day -4, 20 gravid adult (SS104 *glp-4*(bn2)) worms from unstarved plate, were set up to lay eggs at 15°C on 9 cm DM plates. By Day -2, gravid worms were removed from the egg-laying plates. On Day -1, the plate with mixed stage worms was shifted to 24°C to induce sterility. On Day 0, 30 L4 (fourth larval stage) worms were picked for each healthspan assay plate and loaded onto the WormGazer^TM^ for automated assessment of healthspan. The following day marked the first day (Day 1) of adulthood for worms, the experiment was terminated on Day 7, after which the plates were removed and manually checked for quality control (for example contaminations and burrowing). If there is any deviation, the plates were censored from data analysis.

### Healthspan Assay with WormGazer

The WormGazer™ was used to continuously image the worms over a period of seven days. The Petri plates were illuminated from top and imaged by camera below, and the chamber’s temperature and moisture content were controlled. The objects moving above the threshold of 10 µm/s were used to classify movement and the mobility endpoints were calculated by the WormGazer analysis pipeline (Zavagno et al. 2024). We used five mobility endpoints: fraction of moving worms (fraction moving); mean speed of moving worms; mean speed of all worms (population-averaged speed, with non-movers contributing zero and therefore equal to fraction moving × mean speed of moving worms); mean hours moving, defined as area under the fraction-moving curve; and mean distance moved, defined as area under the mean-speed-of-all-worms curve.

### RNA Extraction and Sequencing

Following treatment, worms were harvested on Day 3 of adulthood, washed twice in cold M9 buffer (centrifugation: 1,000 rpm, 30 sec), and resuspended in 10–20 µL residual buffer. RNA was extracted using QIAzol lysis reagent and freeze thaw method following RNeasy Universal Mini Kit (QIAGEN) manufacturer guidelines, with final elution in RNAse-free water. RNA quality was verified using nanodrop, aliquots of RNA were stored at −80°C, and samples were submitted to Novogene for bulk RNA sequencing. A total of 16 samples were submitted for sequencing with 4 replicates for each treatment condition.

### RNA-seq Data Analysis

Raw data from RNA sequencing were processed by Novogene to remove reads with adapters, reads containing >10% ambiguous nucleotides, and low-quality reads with Phred quality ≤ 5 for over 50% bases. Clean reads were mapped to *C. elegans* reference genome (WBcel235) with HISAT2 (v2.0.5) (D. Kim et al. 2019). Aligned reads were assembled with StringTie (v1.3.3b) (Pertea et al. 2015), and gene-level raw counts were generated with FeatureCounts (v1.5.0-p3) (Liao et al. 2014) and normalized to Fragments Per Kilobase Million (FPKM) values for downstream analysis. Differential expressions for each treatment versus control was assessed with DESeq2 (v1.20.0) (Love et al. 2014), and genes with Benjamini-Hochberg-adjusted p value ≤ 0.05 and |log2 fold-change| ≥ 1 were labelled as differentially expressed genes (DEGs). The heatmaps were generated using pheatmap package (Kolde 2025) in R with log2 transformed FPKM values. Euclidean distance metric for genes and samples, and Ward.D2 hierarchical clustering method were used for the heatmap of transcriptome-wide expression. Gene Ontology (GO) (Ashburner et al. 2000; The Gene Ontology Consortium et al. 2023) biological process and KEGG pathway (Kanehisa et al. 2025) enrichment analysis were performed using clusterProfiler (Wu et al. 2021) in R using a common background (union of expressed genes) and BH adjustment (pAdjustMethod = “BH”, pvalueCutoff = 0.05, qvalueCutoff = 0.05). Longevity associated genes were extracted from the KEGG Longevity regulating pathway for worms (cel04212) and their overlap with the DEGs was analyzed. Differential transcription factor (TF) activity was estimated from the differential expression of the target genes for each treatment versus control using the CelEst R Shiny app (network v1.1, default multivariate-linear-model method) (Perez 2025, 2024). Gene expression data was used with gene id and signed -log_10_(p-value) (positive for up-regulated, negative for down-regulated) for TF activity analysis. TFs with adjusted p-value < 0.1 were reported as significantly activated or repressed. The protein-protein interaction network of DEGs was constructed using STRING database v12.0 (Szklarczyk et al. 2023) and combined score cutoff ≥ 0.7. Network visualization and analysis were performed using Cytoscape (Shannon et al. 2003). UpSet plots (Lex et al. 2014; Conway et al. 2017) were created to visualize the gene set intersections between treatments.

### Module-Focused Gene Panel and Block-Structured Heatmap

To summarize recurring signals from enrichment analysis, we defined four modules using enriched GO/KEGG term sets identified in this study (glutathione/detoxification, sphingolipid/lipid remodeling, innate immunity, and longevity/stress; exact terms IDs are listed in Supplementary Table S13). For each contrast (Curcumin vs Control, Sulforaphane vs Control, Combination vs Control), DEGs filtered at *padj* ≤ 0.05 and |log2FC| ≥ 1 and merged by gene. For each module, we took the union of genes annotated to its GO/KEGG term IDs across all the three contrasts (using clusterProfiler results for both up- and down-regulated sets), ranked genes within the module by the maximum absolute log2 fold change across the three contrasts, and selected the top 15. Genes appearing in more than one module were assigned to the module where they ranked highest; duplicates were removed to yield the final panel. Expression values for these genes were log2(FPKM + pseudocount), with the pseudocount set to 0.5 times the minimum non-zero FPKM in the dataset. For visualization, values were scaled by row (z-scores) and plotted without clustering in a block-segmented heat map with samples ordered by treatment.

## Results

Aging is associated with a marked decline in mobility (Rantakokko et al. 2013). In humans, lower-extremity function assessed by the Short Physical Performance Battery (SPPB) is associated with all-cause mortality risk (J. M. Guralnik et al. 1994; Jack M. Guralnik et al. 2000). Similarly, *C. elegans* exhibit a significant reduction in mobility with advancing age (Huang et al. 2004). In this study, we therefore used automated whole-organism phenotyping and quantified five mobility endpoints: fraction of moving worms (fraction moving); mean speed of moving worms; mean speed of all worms (population-averaged speed, non-movers counted as zero); and two cumulative (AUC) summaries, hours moved (area under the fraction-moving curve) and distance moved (area under the mean-speed-of-all-worms curve). Using these endpoints, we screened the selected natural products/plant extracts (berberine, bergamot extract, curcumin, ginkgo, liquorice root extract, *Rhodiola rosea* root extract, *Salvia miltiorrhiza* root extract, and sulforaphane) and the nutritional supplement nicotinamide riboside (**Figure 1a**).

**Figure 1:**
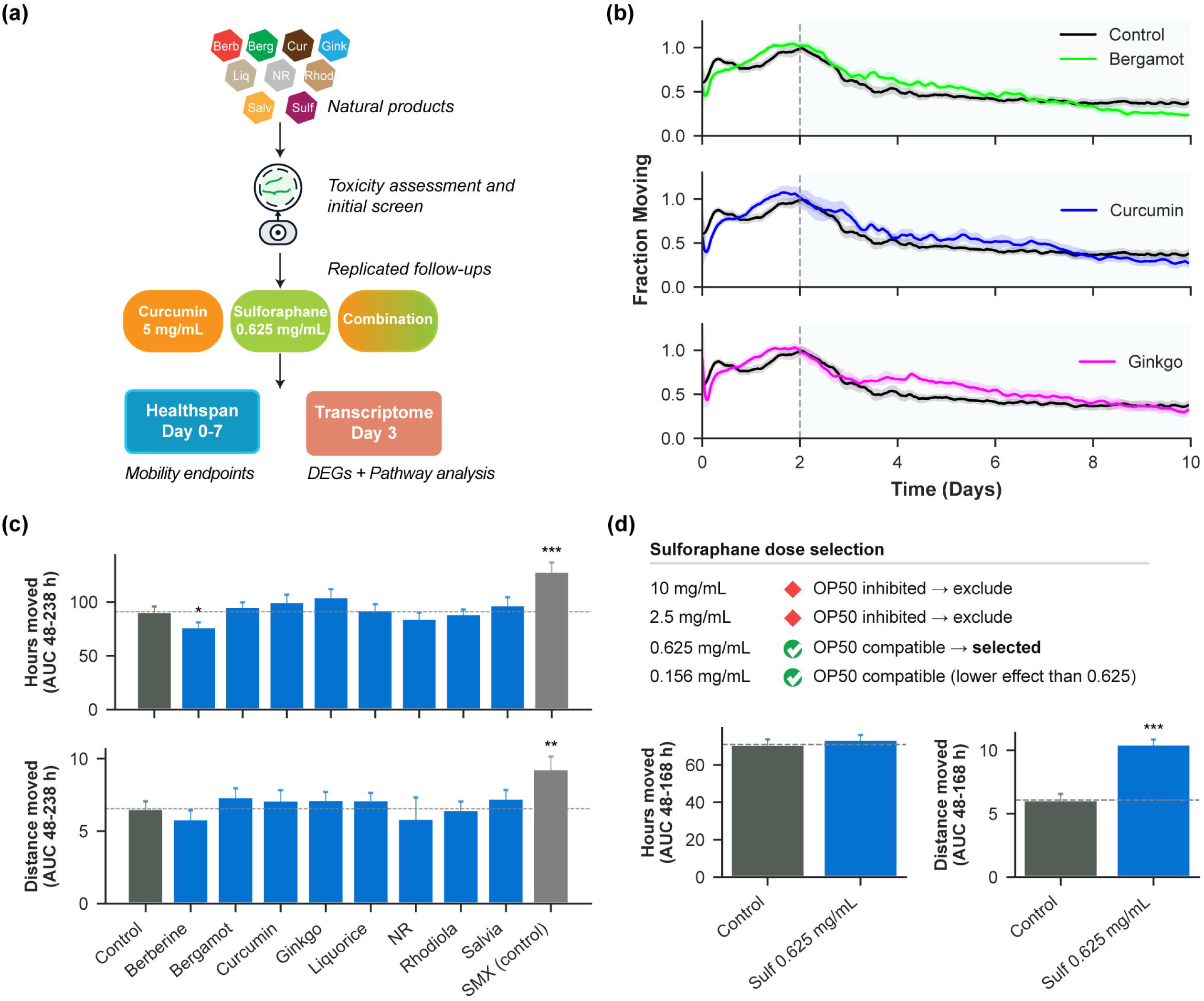
Study overview, initial screen, and sulforaphane dose selection. (a) Schematic workflow of the study’s initial screen (non-replicated) of the natural products/plant extract/nutraceutical followed by selection and replicated follow-up investigation (curcumin 5 mg/mL, sulforaphane 0.625 mg/mL, combination at same concentration), and Day-3 transcriptome profiling (DEGs, gene/pathway enrichment, TF activity inference, protein-protein interaction networks). (b) Representative screen fraction-moving time courses. The dashed line marks Day 2; the shaded region indicates the post-Day 2 interval used for AUC summaries. (c) Screen cumulative summaries (AUC 48-238 hours) for hours moved and distance moved (mean ± SEM from technical replicates). Sulforaphane is excluded here due to bacterial inhibition at screen concentration; see panel (d). SMX is a platform assay control and is not used for head-to-head comparison with interventions. (d) Sulforaphane dose selection in the screen: bacterial (OP50) growth was inhibited at 2.5 and 10 mg/mL, but not at ≤ 0.625 mg/mL. Screen cumulative summaries (AUC 48–168 hours) for control vs sulforaphane 0.625 mg/mL are shown for hours moved and distance moved (mean ± SEM). Stars denote a one-sided test on ΔAUC (treatment vs matched control): * p < 0.05, ** p < 0.01, *** p < 0.002; NS otherwise.

### Qualitative Screening Identified No Obvious Gross Developmental or Reproductive Defects

We first performed a qualitative pre-screen to assess whether any intervention caused obvious gross developmental or reproductive impairment at the concentrations used for the initial mobility screen. Across the tested concentrations (10, 5, and 2.5 mg/mL), none of the compounds produced obvious gross defects in worm growth or progeny production in this qualitative assessment. We therefore used 10 mg/mL for the initial screen for all compounds except sulforaphane. Sulforaphane was treated separately because concentrations ≥ 2.5 mg/mL visibly reduced OP50 lawn density, raising a potential dietary-restriction confound rather than a direct worm-specific effect. An OD600-based bacterial growth assay confirmed inhibition at 2.5 and 10 mg/mL but not at concentrations ≤ 0.625 mg/mL. We therefore used 0.625 mg/mL sulforaphane in subsequent healthspan assays (**Figure 1d; Supplementary Figure S1**).

### Natural Product Interventions Rescue Age-Related Decline in Mobility

We conducted an exploratory initial screen without independent biological replicates to evaluate the effects of the selected interventions on *C. elegans* healthspan (Methods; **Figure 1a**). Sulfonamide antibiotic sulfamethoxazole (SMX, 16 µg/mL), a compound known to extend healthspan and lifespan in *C. elegans* via inhibition of folate synthesis in bacteria, was used as an assay control, and was not used for head-to-head comparison with test compounds (Liu et al. 2013; Virk et al. 2012; Zavagno et al. 2024). Consistent with previous studies (Zavagno et al. 2024; Collins et al. 2018; Glenn et al. 2004), we observed a gradual age-associated mobility decline, with mobility peaking at Day 2 of adulthood and declining thereafter in control worms. Among the interventions, bergamot, curcumin, ginkgo, salvia and sulforaphane showed post-Day 2 improvements in one or more mobility endpoints (**Figure 1b**, **Supplementary Figure S2–S5)**. Using the cumulative summaries: hours moved and distance moved, these five treatments exceeded control, with sulforaphane showing the largest increase in distance moved in this exploratory initial screen (**Figure 1c-d; Supplementary Figure S6a–b**).

Although several interventions improved late-adult mobility in the screen, we advanced sulforaphane and curcumin because they showed strong and distinct patterns in the late-adult mobility endpoint gains. Sulforaphane showed the largest increases in speed-based endpoints (speed of moving worms, speed of all worms, distance moved), whereas curcumin most consistently increased time active (fraction moving, hours moved). Ginkgo performed closely to curcumin on the cumulative summaries (**Figure 1c**). Thus, we evaluated curcumin (5 mg/mL), sulforaphane (0.625 mg/mL), and their combination in two independent biological replicates.

### Effects of Curcumin on Mobility

Treatment with curcumin alone influenced several mobility endpoints throughout the observation period. It transiently increased the fraction of moving worms between Days 1–2 and more consistently from ∼Day 3.5–7, relative to the age-related decline observed in control worms (**Figures 2a; Supplementary Figure S7a**). For speed endpoints, curcumin treatment initially reduced the speed of moving worms (mean speed among movers) through ∼Day 4, followed by a sustained increase thereafter (**Figure 2b; Supplementary Figure S7b**). The speed of all worms (population-averaged speed) was variable early but improved from ∼Day 3.5 onward (**Figure 2c; Supplementary Figure S7c**). Correspondingly, hours moved (AUC Days 2–7) was significantly increased in both independent experiments, whereas effects in the early window (Day 0–2) were inconsistent across replicates. Despite later speed gains, the cumulative summary endpoint, distance moved (area under the mean-speed-of-all-worms, Days 2–7) did not reach significance with curcumin treatment (**Figure 2d; Supplementary Figure S7d, S8**).

**Figure 2:**
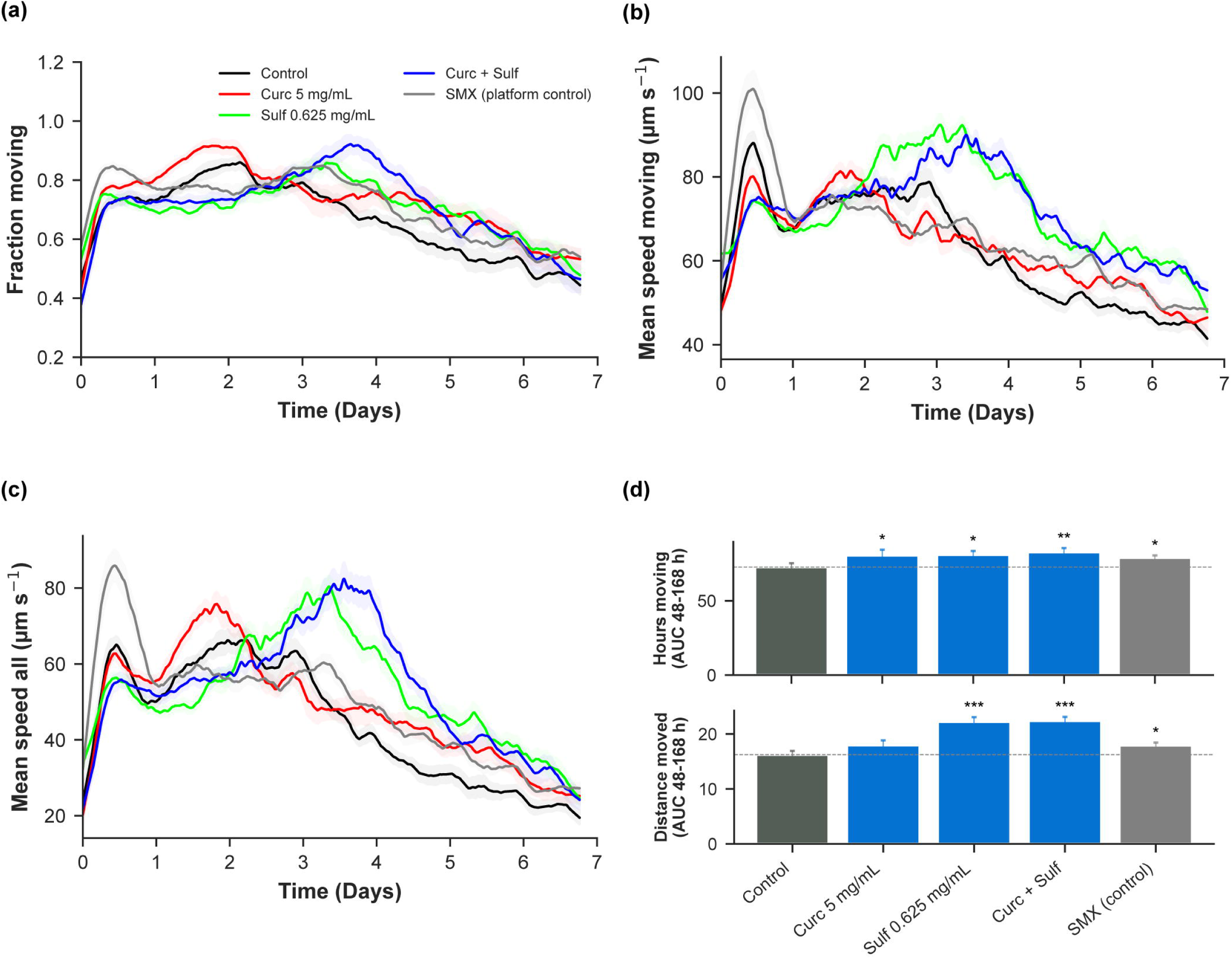
Mobility under curcumin (5 mg/mL), sulforaphane (0.625 mg/mL), and their combination. (a–c) Time courses for fraction moving, mean speed of moving, and mean speed of all worms (Replicate 1; mean ± SEM shown). SMX is a platform control. (d) cumulative summaries (AUC 48–168 h) for hours moved and distance moved relative to matched controls. Stars denote a one-sided test on ΔAUC (treatment vs matched control): * p < 0.05, ** p < 0.01, *** p < 0.002; NS otherwise. Replicate 2 is shown in Supplementary Figure S7 (identical layout).

### Effects of Sulforaphane on Mobility

Sulforaphane treatment revealed a distinct temporal pattern. Relative to control, sulforaphane reduced the fraction of moving worms between Days 0**–**2 but increased it from **∼**Day 3**–**7 in both replicates (**Figures 2a; Supplementary Figure S7a**). For the speed endpoints, sulforaphane reduced both the speed of moving worms (mean speed among movers) and the speed of all worms (population-averaged speed) early in the assay, followed by a sustained increase in both measures after Day 2 (**Figures 2b–c; Supplementary Figures S7b–c**). This post-Day 2 enhancement was often the largest among treatments, with sulforaphane outperforming curcumin in this interval. Mirroring the above pattern, hours moved (area under the fraction-moving curve Days 2–7) increased significantly versus control in both replicates, whereas in the Day 0–2 window it decreased. Consistent with the speed gains, distance moved (area under the mean-speed-of-all-worms Days 2–7) increased significantly versus control, with one replicate showing the largest gain among treatments (**Figures 2d; Supplementary Figures S7d, S8**).

### Combined Effect of Curcumin and Sulforaphane on Mobility Profile

The combined treatment of curcumin and sulforaphane improved the fraction of moving worms from ∼Day 2.5–6.5, although the magnitude of this effect varied between replicates (**Figures 2a; Supplementary Figure S7a**). For the speed endpoints, the combination reduced the speed of moving worms during Day 0–2, then increased it after ∼Day 3, with relative magnitude versus sulforaphane differing by replicate (**Figure 2b; Supplementary Figure S7b**). In both replicates, during Days 2–3.5, the combined treatment effect was lower than that of sulforaphane alone, but from ∼Day 3.5 onward, the effects became similar. In replicate 1, this similarity persisted until Day 7, whereas in replicate 2, it lasted until Day 5.5, after which the effect of the combination declined relative to sulforaphane. The mean speed of all worms followed a similar pattern, with the combination showing a positive effect from Day 2.5 onwards (**Figure 2c; Supplementary Figure S7c**). Following the effect on fraction moving, hours moved (AUC, Days 2–7) was significantly increased versus control in both replicates, whereas in the Day 0–2 window it decreased. Reflecting the combined treatment impact on speed, distance moved (AUC Days 2–7) was significantly increased in both replicates, generally exceeding curcumin alone and being similar to sulforaphane (**Figures 2d; Supplementary Figures S7d, S8**). Overall, the combination treatment produced late-adult mobility gains comparable to sulforaphane across hours moved and distance moved.

### Curcumin and Sulforaphane Show Distinct Transcriptional Responses, with Broader Remodeling in the Combination

To define the transcriptional responses to curcumin and sulforaphane, we performed RNA-seq on Day 3 adult worms treated with curcumin (5 mg/mL), sulforaphane (0.625 mg/mL), their combination at the same concentrations, or control, with four independent biological replicates per condition (**Figure 3a; Supplementary Table S1**). A transcriptome-wide hierarchical clustering analysis (Euclidean distance, ward.D2; **Supplementary Figure S9**) showed that replicates clustered by treatment and separated across conditions.

**Figure 3:**
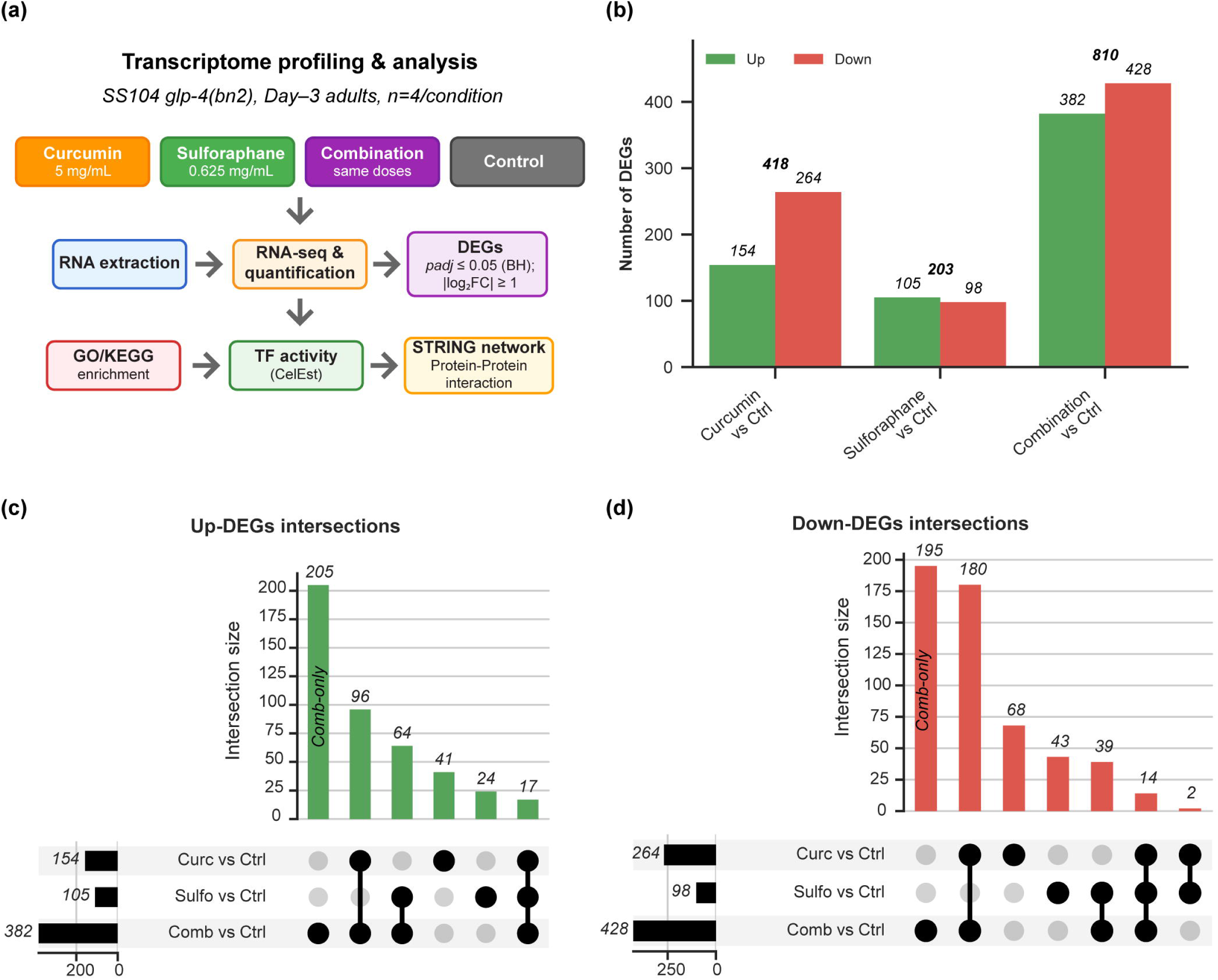
RNA-seq design and DEG landscape. (a) Transcriptome profiling & analysis: SS104 *glp-4*(bn2), Day-3 adults, n = 4/condition; downstream analyses include DEGs, GO/KEGG enrichment, TF activity inference (CelEst), and STRING protein-protein interaction networks. (b) Numbers of Up and Down DEGs (*padj* ≤ 0.05; |log FC| ≥ 1) for each treatment; totals above pairs: Curcumin 154/264 (418), Sulforaphane 105/98 (203), Combination 382/428 (810). (c) UpSet plot of Up-DEGs across Curc, Sulf and Comb (vs Ctrl). Set-size bars show totals; the Comb-only intersection (205 genes) is highlighted. (d) UpSet plot of Down-DEGs across Curc, Sulf and Comb (vs Ctrl). Totals shown; the Comb-only intersection (195 genes) is highlighted.

Using a significance threshold of *padj* ≤ 0.05 and |log2FC| ≥ 1, the combination treatment produced the largest differential expression response, with 382 up-regulated and 428 down-regulated genes, compared with 154 up / 264 down for curcumin and 105 up / 98 down for sulforaphane (**Figure 3b; Supplementary Tables S2-S4**). Treatments also differed substantially from one another: pairwise comparisons identified 643 DEGs between curcumin and sulforaphane, 218 between curcumin and the combination, and 569 between sulforaphane and the combination (**Supplementary Figure S10**), indicating that the combination produces a broader transcriptional profile that remains distinct from either single-compound treatment.

We next compared the curcumin- and sulforaphane-responsive DEG sets relative to control. The two treatments showed limited same-direction overlap, with only 17 co-up-regulated and 16 co-down-regulated genes. In addition, 15 genes were regulated in opposite directions, including ten transcripts such as *cyp-14A3*, *cyp-35A3*, and the C-type lectin genes *clec-60* and *clec-10* that were induced by curcumin but suppressed by sulforaphane, whereas five transcripts showed the opposite pattern (**Supplementary Table S5**). Together, these patterns, along with the large direct difference between curcumin and sulforaphane, support the conclusion that the two compounds elicit distinct transcriptional programs.

To benchmark our data, we compared sulforaphane-responsive genes from our study with those reported by Sedore *et al*. (100 µM, N2 strain) (Sedore et al. 2025). Despite lower dose, strain difference and time point variation, 38% of their Day 4 up-regulated genes (11 out of 29) and 12% of their Day 8 up-regulated genes (45 out of 369) overlapped with ours, with no direction mismatches **(Supplementary Table S6)**. This concordance supports the consistency of our transcriptomics dataset. Comparable curcumin transcriptomes were not available for validation.

### Combination Treatment Broadens Detoxification and Retunes Immune Response

Notably, 400 of the 810 DEGs detected in the combination treatment were combination-specific, 205 uniquely up-regulated and 195 uniquely down-regulated genes not observed in either single-compound treatment (**Figure 3c,d**). The uniquely up-regulated gene set included additional detoxification-associated genes including canonical stress-response gene *gst-4*, and members of the *ugt* and *cyp* families, whereas the uniquely down-regulated set included 13 additional *clec* genes together with *lys-7*. The combination specific down-regulated genes also included marked repression of *pqm-1* (log2FC = -1.25; *padj* = 5.01e-55), consistent with reduced PQM-1 associated transcription. Together, these results indicate that the combination does not simply recapitulate the single-compound treatment responses but adds a substantial combination-specific component that broadens detoxification-associated transcriptional signatures while extending repression of innate immune effector genes.

### Combination Treatment Retains Single-Compound Pathway Signatures and Broadens Longevity-Associated Programs

We next examined direction-specific GO biological process (BP) enrichment for each treatment to compare curcumin, sulforaphane, and the combination. Curcumin up-DEGs were enriched for membrane lipid remodeling, particularly sphingolipid- and ceramide-related processes together with phospholipid catabolism, whereas curcumin down-DEGs were enriched for host-microbe interaction and defense terms (**Figures 4a,b; Supplementary Table S9**). Sulforaphane up-DEGs prominently featured glutathione and sulfur-compound metabolism, with additional enrichment for response-to-organism and immune/defense-related categories. In contrast, sulforaphane down-DEGs showed a narrower set of defense- and xenobiotic-response terms. The combination treatment retained key features of both single-compound treatment responses, combining lipid-remodeling and glutathione/sulfur-compound signatures in the up-DEGs while showing broad enrichment of immune/defense terms among down-DEGs (**Figures 4a,b; Supplementary Table S9**).

**Figure 4:**
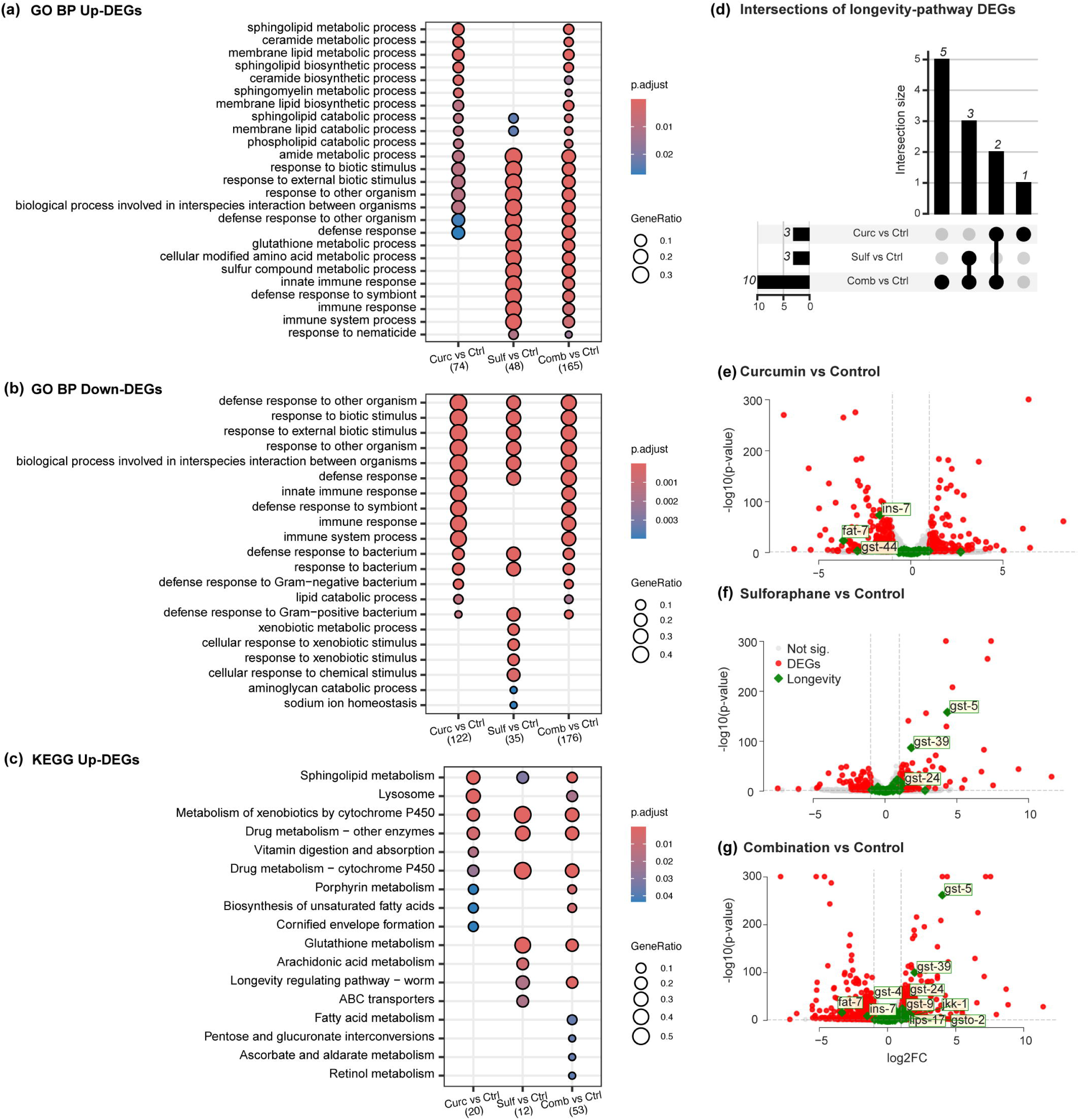
(a) GO Biological Process (Up-DEGs), (b) GO Biological Process (Down-DEGs), and (c) KEGG pathways (Up-DEGs) for Curc (curcumin), Sulf (sulforaphane), and Comb (curcumin+sulforaphane) versus control. Dot size indicates GeneRatio; dot color indicates BH-adjusted p (*padj*). Numbers in parentheses denote the annotated input used (mapped genes). (d) UpSet plot of KEGG Longevity regulating pathway-worm (cel04212) genes that are differentially expressed vs control in each treatment (Curc = curcumin; Sulf = sulforaphane; Comb = curcumin+sulforaphane). Counts reflect the direct overlap of the full pathway gene set with treatment DEGs (*padj* ≤ 0.05; |log2FC| ≥ 1); bars show intersection sizes (the Comb-only set is largest). (e–g) Volcano plots for Curc, Sulf, and Comb highlighting longevity-pathway genes (green diamonds). Dashed vertical lines mark |log2FC| = 1; the y-axis shows −log10(p). Selected significant genes are labeled: Curc:*ins-7*, *fat-7*, *gst-44* (down); Sulf:*gst-5*, *gst-39*, *gst-24* (up); Combo:*gst-5, gst-39, gst-4, gst-24,* gs*t-9, jkk-1, gsto-2, lips-17* (up), with *ins-7* and *fat-7* remaining suppressed.

At the KEGG pathway level, xenobiotic- and drug-metabolism modules were enriched for all three treatments, whereas glutathione metabolism was enriched only for sulforaphane and the combination (**Figure 4c; Supplementary Table S10**). Curcumin up-DEGs additionally showed strong enrichment for membrane-lipid and lysosomal pathways, with lysosome enriched for curcumin and the combination but not for sulforaphane alone. Importantly, Longevity regulating pathway-worm was enriched in the up-DEGs of sulforaphane and the combination, but not curcumin, and the combination contributed the largest treatment-specific subset within this pathway (**Figures 4c,d; Supplementary Table S10**). Within this longevity-pathway overlap, sulforaphane up-regulated *gst-5*, *gst-39*, and *gst-24*, consistent with a glutathione-linked detoxification signature, whereas curcumin down-regulated *ins-7* (an IIS agonist), *fat-7* (a fatty-acid desaturase), and *gst-44*. The combination retained these single-compound treatment features while additionally inducing *gst-4*, *gst-9*, *gsto-2*, *jkk-1* (a JNK-pathway kinase), and *lips-17* (a lysosomal lipase). *ins-7* and *fat-7* remained suppressed, whereas *gst-44* was no longer significantly altered (**Figures 4e,f**). Together, these results indicate that the combination preserves the major pathway features of each single-compound treatment while broadening engagement of detoxification- and longevity-associated programs.

### Combination Treatment Integrates Key Transcription Factor Programs

We inferred transcription factor (TF) activity using the CelEst multivariate linear model (*padj* < 0.1; Methods) (Perez 2025). Curcumin was associated with inferred repression of the tubby-like factor TLP-1 and activation of the zinc-finger TF ZTF-22, whereas sulforaphane was associated with inferred activation of SKN-1, several nuclear hormone receptors (NHR-20, NHR-70, NHR-77, NHR-91), SMA-3, ATHP-1 and MEMI-2. The combination retained key sulforaphane-linked signals, including SKN-1, NHR-20, and NHR-70, while also showing inferred activation of ATFS-1, SMA-3, LIN-40, and FKH-2, together with inferred repression of TLP-1 and GMEB-1 (**Figures 5a-c, Supplementary Table S11).** Overall, the combination TF profile is consistent with retention of detoxification- and lipid-remodeling-associated regulatory features, in line with the gene- and pathway-level signatures described above. Representative DEGs in the combination that align with these inferred TF programs include *gst-4* for the detoxification axis and *lips-17* for the lipid-remodeling axis.

**Figure 5:**
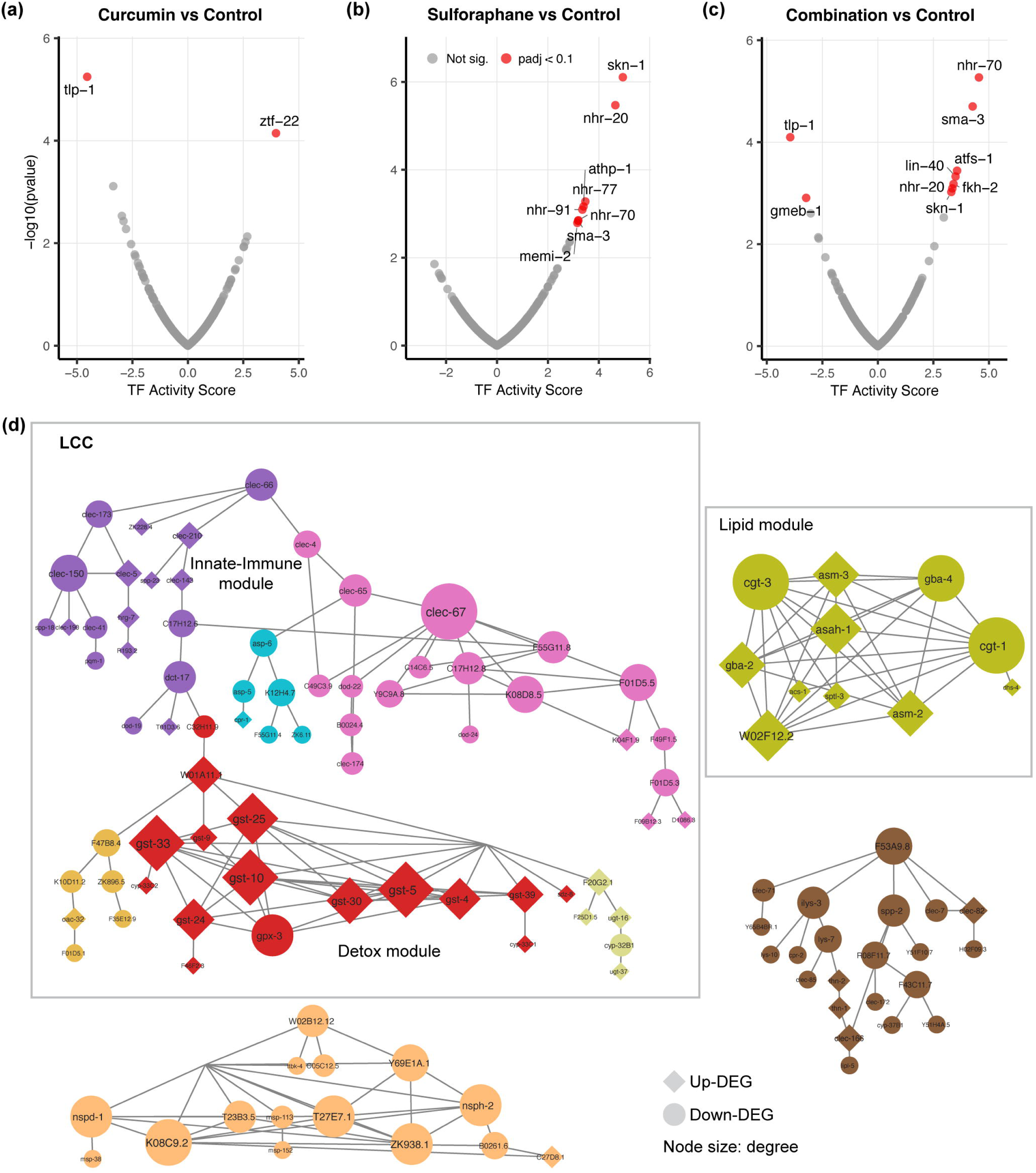
Transcription-factor activity inference (CelEst). TF activity score (x) versus significance (y = −log10(pvalue)) for (a) Curc, (b) Sulf, and (c) Comb vs control. Points in red indicate *padj* < 0.1; gray = not significant. (d) STRING protein-protein network of DEGs under Combination treatment, showing the four largest connected components. Edges: STRING v12.0, combined score ≥ 0.7; nodes are DEGs only (*padj* ≤ 0.05; |log2FC| ≥ 1); isolates removed; communities by Glay (node color = community). The largest connected component (LCC; 71 genes) co-locates a detox GST cluster (*gsto-2, gst-4, -5, -9, -10, -24, -25, -30, -33, -39* up, diamonds) with innate-immune CLECs (mostly down, circles). A distinct lipid module (11 genes) is retained (asah-1 up; cgt-1, -3 Down). The complete network is shown in Supplementary Figure S13.

### Combined Curcumin and Sulforaphane Expand and Integrate Detoxification and Immune Networks

We next examined whether the pathway-level signatures identified above were also reflected at the network level. Under curcumin, the protein-protein interaction (PPI) network comprised 113 nodes across 24 connected components. Its largest connected component (LCC; 43 genes) was predominantly down-regulated (34 down, 9 up) and was enriched for innate-immune and defense-associated genes, including seven *clec* genes (5 down, 2 up) and three *irg* genes (2 down, 1 up); *clec-67* (down) and *irg-4* (up) were the top-degree nodes (degree 7 each). A separate, highly connected eight-node lipid module captured the curcumin lipid-remodeling signal (*asah-1* up; *cgt-1* and *cgt-3* down) (**Supplementary Figure S11; Supplementary Table S12**).

Sulforaphane yielded a smaller 40-node network whose LCC (11 genes) was centered on detoxification genes, including *gst-5*, *gst-33*, *gst-24*, *gst-30*, and *gst-39* (all up-regulated), alongside a small defense-associated mini-cluster containing *clec-52*, *clec-60*, and *ilys-3* (all down-regulated) (**Supplementary Figure S12; Supplementary Table S12**).

In the combination treatment, the PPI network expanded to 214 nodes across 37 connected components (**Supplementary Figure S13; Supplementary Table S12**). To improve readability, **Figure 5d** shows the four largest connected components of this network, while Supplementary Figure S13 shows the complete network. Notably, the detoxification and innate-immune modules were co-located within the same LCC (71 genes). This LCC contained 10 *gst*/*gsto* genes, all up-regulated (*gsto-2, gst-4, gst-5, gst-9, gst-10, gst-24, gst-25, gst-30, gst-33, gst-39 up*), and 12 *clec* genes that were predominantly down-regulated (*clec-4, clec-41, clec-65, clec-66, clec-67, clec-150, clec-173, clec-174* down *and clec-5, clec-143, clec-190, clec-210* up). In contrast, the lipid-remodeling signal remained as a distinct highly connected 11-node module, with *asah-1* up-regulated and *cgt-1* and *cgt-3* down-regulated (**Figure 5d**).

Together, these network analyses indicate that the combination integrates the detoxification and innate-immune signatures into a shared connected framework, while retaining the curcumin-associated lipid module as a separate network component. This pattern is consistent with a broader, coordinated transcriptional response under the combination treatment.

### Replicate-Level Visualization of Key Aging-Related Modules Supports the Combination Signature

Guided by the transcriptomic results above (GO/KEGG enrichment, TF-activity inference, and protein-protein interaction networks), we focused on four aging-related modules that repeatedly emerged: glutathione/detoxification, sphingolipid/lipid remodeling, innate immunity, and longevity/stress. To visualize these signals across individual replicates, we generated a block-segmented heatmap of curated genes drawn from enriched GO/KEGG term sets, ranked within each module by maximum absolute log2 fold change across contrasts, and deduplicated as described in Methods (**Figure 6; Supplementary Table S13**).

**Figure 6:**
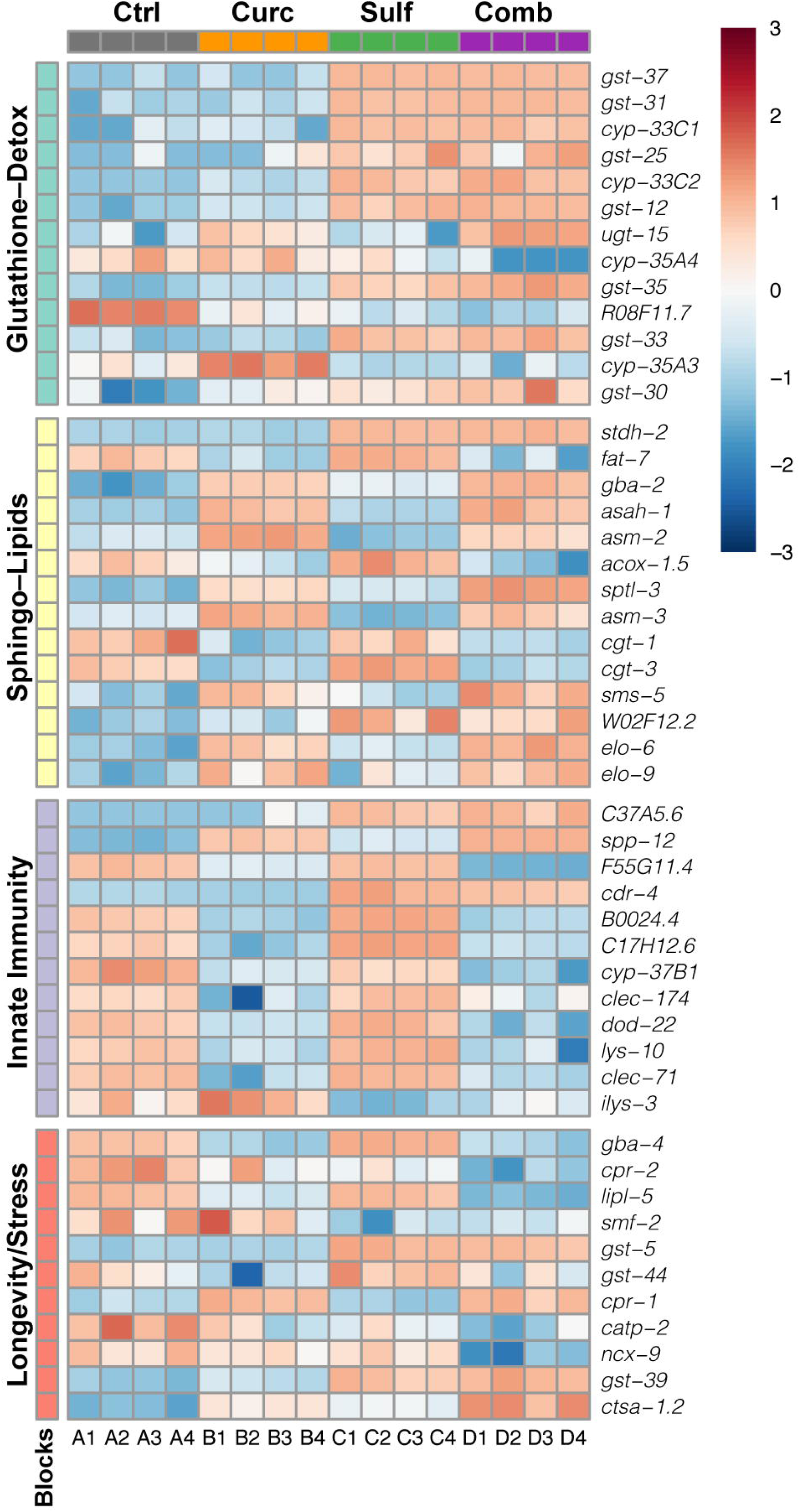
Block-segmented heatmap of curated genes from key aging modules. Rows are 50 differentially expressed genes selected from enriched GO/KEGG sets and grouped into four modules: glutathione/detox (teal), sphingolipid/lipid (yellow), innate immunity (purple), and longevity/stress (coral). Within each module, genes were ranked by the maximum absolute log2 fold change in curcumin (Curc), sulforaphane (Sulf), or their combination (Comb) versus control (Ctrl); the top 15 per module were retained and genes occurring in multiple modules were assigned to the highest-ranked module (Methods; Supplementary Table S13). Columns show the 16 RNA-seq libraries (A1–A4 = Ctrl, B1–B4 = Curc, C1–C4 = Sulf, D1–D4 = Comb; group annotation bar above). Values are row-wise z-scores of log2-FPKM (diverging scale centered at 0).

In the glutathione/detox block, most genes were low in control and curcumin, whereas sulforaphane showed coherent induction that was largely maintained in the combination. In the sphingolipid/lipid block, curcumin showed a characteristic remodeling pattern, including higher *gba-2, asah-1, asm-2, asm-3* and *sptl-3* together with lower *fat-7, acox-1.5, cgt-1,* and *cgt-3*; most of these features persisted in the combination. In the innate-immune block, several lectin-and lysosome-associated genes, including *clec-174*, *clec-71*, *dod-22*, and *lys-10*, were lower in curcumin and were generally further reduced in the combination, although a subset of secreted or defense-associated genes such as *spp-12* and *cdr-4* showed modest induction. In the longevity/stress block, sulforaphane-associated glutathione genes such as *gst-5* and *gst-39* remained elevated in the combination, whereas *gst-44* remained lower under curcumin and in the combination treatment; additional genes in this block, including *gba-4*, *lipl-5*, *ncx-9,* and *catp-2*, also differed across treatments.

Together, this module-level view is consistent with the combination preserving prominent features of each single-compound treatment, sulforaphane-associated glutathione/detoxification signatures and curcumin-associated lipid remodeling, while extending repression of several innate-immune effectors. This visual summary is consistent with broader late-adult maintenance-oriented transcriptional remodeling under the combination treatment.

## Discussion

Aging is accompanied by progressive decline in physiological and functional capacity that culminates in frailty and loss of resilience. Among measurable functional traits, movement capacity has emerged as a quantifiable biomarker of biological aging that integrates neuromuscular, metabolic, and mitochondrial performance. In humans, gait speed and cardiorespiratory fitness are among the strongest predictors of mortality and morbidity, outperforming chronological age as a prognostic index (Gaesser et al. 2025; Studenski et al. 2011). In *Caenorhabditis elegans*, locomotor decline closely parallels frailty trajectories observed in mammals and can be longitudinally tracked with high temporal precision (Herndon et al. 2002; Huang et al. 2004; Newell Stamper et al. 2018; Tissenbaum 2012). After the Day 2 peak in mobility, worms exhibit progressive loss of motility that correlates strongly with individual survival, supporting locomotion as a surrogate marker for healthspan (Kawamura and Maruyama 2019; Hahm et al. 2015).

Using mobility as a functional healthspan-related readout, we examined how curcumin and sulforaphane influence late-adult locomotor decline in *C. elegans*. Both compounds are well-known hormetic agents that modulate stress-response pathways, reduce inflammation, and improve metabolic homeostasis (M. Cheng et al. 2025; Alves et al. 2025). In follow-up assays, curcumin, sulforaphane, and their combination all improved mobility after Day 2 of adulthood, with sulforaphane producing the strongest late-phase effects on mean speed, distance moved, and hours moved, whereas curcumin showed a distinct benefit profile, with more pronounced effects on time-active measures such as fraction moving and hours moved than on speed-based metrics (**Figure 2**; **Supplementary Figures S7, S8**). The combination largely paralleled sulforaphane at the phenotypic level but was associated with broader transcriptomic remodeling, consistent with the two compounds engaging overlapping yet non-identical molecular programs.

At the molecular level, curcumin and sulforaphane elicited distinct yet partially convergent transcriptional responses that were broadened in the combination treatment. Sulforaphane was associated with induction of multiple detoxification-related genes, including *gst*, *cyp*, and *ugt* family members, consistent with an SKN-1-linked detoxification signature, whereas curcumin more prominently altered genes linked to lipid and sphingolipid remodeling and reduced the expression of several innate immune effectors, including multiple *clec* genes (**Figure 4a-c**). The combination retained major features of both single-compound treatment responses while adding combination-specific changes, including additional detoxification-associated transcripts together with further repression of lectin- and lysozyme-associated genes. Together, these signatures are consistent with a broader maintenance-oriented transcriptional program characterized by enhanced glutathione and xenobiotic metabolism, coordinated lipid remodeling, and reduced expression of several innate immune effectors.

Mechanistically, Nrf2/SKN-1 orchestrates cytoprotective gene expression from nematodes to humans (An and Blackwell 2003). In mammalian systems, sulforaphane activates Nrf2 and promotes glutathione-linked antioxidant and detoxification responses (Suzuki et al. 2023; Alves et al. 2025). Consistent with this framework, our data showed inferred activation of SKN-1 under sulforaphane and the combination, together with induction of multiple glutathione-related genes and combination-specific up-regulation of *gst-4* (**Figure 5b,c**). We also observed inferred activation of NHR-70 and combination-specific induction of *lips-17*, supporting a link between the combination response and lipid-remodeling or lysosome-associated programs. In addition, the combination altered other regulatory nodes, including up-regulation of *jkk-1* and repression of *pqm-1*, further supporting the view that the combination response extends beyond glutathione detoxification to additional maintenance-associated programs.

Curcumin has been reported to modulate inflammatory signaling and lipid metabolism in mammalian systems, including attenuation of MAPK/NF-κB-associated responses (Lee et al. 2023; M. Cheng et al. 2025). Consistent with this broader context, curcumin in our study was associated with reduced expression of several innate immune effectors, including *lys-4* and multiple *clec* genes, together with transcriptional changes linked to sphingolipid, ceramide, and phospholipid remodeling. In *C. elegans*, sphingolipid and ceramide metabolism, together with autophagy-linked lipid turnover, have established roles in aging-related physiology and longevity (Lapierre et al. 2011; Cutler et al. 2014). These signatures suggest that curcumin engages a distinct maintenance-associated program characterized by lipid remodeling alongside lower expression of selected immune effectors, consistent with reduced basal immune activation and broadly aligned with the idea that age-associated inflammatory signaling contributes to functional decline (Franceschi and Campisi 2014). These signatures provide a plausible molecular context for the distinct late-adult mobility benefits observed with curcumin.

Cross-species studies support the biological relevance of the detoxification- and immune-modulatory signatures observed here. In mammals, sulforaphane induces Nrf2-dependent cytoprotective programs in tissues such as liver, intestine, and brain, enhancing resilience to oxidative and inflammatory stress (Dinkova-Kostova et al. 2017; Clarke et al. 2011; Egner et al. 2014). Curcumin similarly modulates inflammatory signaling in mammalian cells, including repression of NF-κB associated responses and reduced cytokine output (Y. Xu and Liu 2017; Y. S. Kim et al. 2007; Karimian et al. 2017). More broadly, metabolite-linked stress responses have also been implicated in conserved aging-related pathways, as illustrated by β-hydroxybutyrate in nematodes and mammals (Newman and Verdin 2017; Edwards et al. 2014; Shimazu et al. 2013). Together, these parallels place our findings within a broader framework of conserved stress- and maintenance-associated responses.

Mobility provides an integrative functional readout of organismal state, reflecting the combined influence of neuromuscular, metabolic, and cellular maintenance processes. In our assays, improved late-adult mobility was accompanied not by a discrete locomotion-associated transcriptional signature, but by broader changes in detoxification, lipid remodeling, and innate immune effector expression. This pattern is consistent with the possibility that curcumin and sulforaphane preserve movement indirectly by shifting the animal toward a more maintenance-oriented molecular state, rather than by directly inducing a specific mobility program. In this context, the combination treatment is notable for coupling sulforaphane-associated detoxification signatures with curcumin-associated lipid-remodeling and immune-effector changes, while remaining functionally similar to sulforaphane at the behavioral level. Taken together, these data suggest that preserved late-adult mobility may reflect coordinated maintenance-related programs spanning detoxification, lipid remodeling, and innate immune effector modulation rather than a discrete locomotion-specific transcriptional response. Future experiments will be needed to determine which of these molecular programs are causally linked to the observed mobility phenotype.

The cross-species relevance of the pathways highlighted here suggests that these findings may have broader biological significance beyond *C. elegans*. Sulforaphane can reach physiologically relevant plasma concentrations through cruciferous vegetable intake, and curcumin has an established safety record in humans (Egner et al. 2011; A. L. Cheng et al. 2001; Lao et al. 2006). Both compounds engage conserved stress- and maintenance-associated pathways, supporting their further evaluation in human healthspan-related contexts. Future studies should define optimal dosing and timing and determine whether the molecular signatures observed here translate into measurable benefits in humans, particularly in tissues with high oxidative burden such as muscle, liver, and brain. Beyond individual agents, our data support the rationale for multi-target nutraceutical strategies that combine compounds with partially distinct but complementary hormetic signatures. Cross-species transcriptomic comparisons may help identify conserved modules associated with improved functional outcomes and guide more rational combination design. Pairing such strategies with metabolic co-interventions, including β-hydroxybutyrate supplementation or targeted sphingolipid modulation, represents an additional hypothesis-generating direction that warrants prospective dose–time studies and formal interaction testing.

Finally, mobility can serve as a useful functional endpoint in translational studies. As in *C. elegans*, where age-related movement decline tracks functional state, longitudinal measures such as gait speed and physical activity in humans may provide sensitive readouts of intervention effects (Huang et al. 2004; Hahm et al. 2015; Studenski et al. 2011; Gaesser et al. 2025). Pairing such physiological measures with molecular readouts of stress-response, metabolic, and inflammatory pathways could help define healthspan across scales. More broadly, these findings are consistent with the view that aging reflects progressive loss of coordination across interdependent biological processes rather than deterioration of a single pathway, a systems-level perspective advanced in theoretical studies of aging networks (Vural et al. 2014). In this context, our findings support a systems-level view in which late-adult functional maintenance may reflect coordinated changes across detoxification, lipid remodeling, and innate immune effector programs rather than modulation of a single pathway. These findings suggest that dietary molecules can engage evolutionarily conserved stress- and maintenance-associated pathways linked to preserved late-adult function, offering a conceptual framework for future translational geroscience.

### Conclusions

In *C. elegans*, curcumin and sulforaphane improved late-adult mobility and, in combination, produced a broader module-level transcriptomic response spanning glutathione-linked detoxification, lipid remodeling, tonic down-tuning of several constitutive innate immune effectors, and components of longevity/stress-associated pathways. Functionally, the combination performed similarly to sulforaphane, while molecular analyses (GO/KEGG, TF-activity inference, protein-protein association networks, and replicate-level module visualization) revealed a broadened response with combination-specific additions. Together, these findings support the idea that combinatorial interventions can expand maintenance-associated pathways, even when behavioral readouts approach a ceiling, and provide a basis for future causal tests (dose–response/interaction modeling; *skn-1*, *nhr-70*, *pqm-1* perturbations) and translation to other contexts and species.

## Acknowledgments

The authors acknowledge funding for this work from Thorne and startup funds to N.D.P at the Buck Institute.

## Author Contributions

Conceptualization: R.P.V.A., D.S., K.M., D.W., S.M.P., N.D.P. Methodology: R.P.V.A., D.S., K.M., C.S., D.W., S.M.P., N.D.P. Investigation: S.M., K.M., C.S., D.W. Formal Analysis: R.P.V.A. Data Curation: R.P.V.A. Visualization: R.P.V.A. Resources: N.D.P., S.M.P. Supervision: N.D.P. Writing – Original Draft: R.P.V.A., D.S. Writing – Review & Editing: All authors.

## Data Availability Statement

RNA-seq data have been deposited in GEO under accession GSE326255. All other data are provided in Supplementary Tables in this study.

## SUPPLEMENTARY INFORMATION

Supplemental Information consists of Supplementary Figures S1-S13 and Supplementary Tables S1-S13.

### Supplementary Figures

**Figure S1:** OP50 growth under sulforaphane for dose selection (48 h). Bars show mean ± SD cell counts (cells ×10), n = 8 technical replicates per condition. One-way ANOVA with Dunnett’s adjusted comparisons vs water control: only 2.5 and 10 mg/mL sulforaphane significantly reduced OP50 counts (adjusted p < 0.0001); doses ≤ 0.625 mg/mL were not significant. Accordingly, 0.625 mg/mL was used as the OP50-compatible concentration in worm assays.

**Figure S2:** Fraction-moving trajectories for eight interventions and SMX (platform control), each overlaid with its matched control (gray). Curves span 0–10 days; AUC summaries over the post-Day 2 interval (48–238 h) are reported in the main text (Fig. 1c). SMX is shown to document assay sensitivity and is not used for head-to-head ranking.

**Figure S3:** Speed of moving trajectories for eight interventions and SMX (platform control), each overlaid with its matched control (gray). Curves span 0–10 days; AUC summaries over the post-Day 2 interval (48–238 h) are reported in the main text (Fig. 1c). SMX is shown to document assay sensitivity and is not used for head-to-head ranking.

**Figure S4:** Speed of all worms trajectories for eight interventions and SMX (platform control), each overlaid with its matched control (gray). Curves span 0–10 days; AUC summaries over the post-Day 2 interval (48–238 h) are reported in the main text (Fig. 1c). SMX is shown to document assay sensitivity and is not used for head-to-head ranking.

**Figure S5:** Sulforaphane dose comparison in the screen. (a) Fraction moving, (b) speed of moving (µm·s ¹), and (c) mean speed of all worms (µm·s ¹) for control and sulforaphane at 0.156, 0.625, and 2.5 mg/mL.

**Figure S6:** Sulforaphane dose comparison in the screen: AUC endpoints (48–168 hours). (a) Hours moved (AUC of fraction moving) and (b) distance moved (AUC of mean speed of all worms) for sulforaphane 0.156, 0.625, and 2.5 mg/mL versus the matched control (bars = mean ± SEM from technical replicates). Stars denote a one-sided test on ΔAUC (treatment vs matched control): * p < 0.05, ** p < 0.01, *** p < 0.002; NS otherwise.

**Figure S7:** Mobility under curcumin (5 mg/mL), sulforaphane (0.625 mg/mL), and their combination. (a–c) Time courses for fraction moving, mean speed of moving, and mean speed of all worms (Replicate 2; mean ± SEM shown). SMX is a platform control. (d) AUC (48–168 h) for hours moved and distance moved relative to matched controls. Stars denote a one-sided test on ΔAUC (treatment vs matched control): * p < 0.05, ** p < 0.01, *** p < 0.002; NS otherwise. Replicate 1 is shown in Figure 2 (identical layout).

**Figure S8:** Early-window AUC (Day 0–2). (a) Replicate 1 and (b) Replicate 2. Bars show hours moved (top; AUC 0–48 h) and distance moved (bottom; AUC 0–48 h) for Control, Curcumin 5 mg/mL, Sulforaphane 0.625 mg/mL, Combination, and SMX (platform control). Values are mean ± SEM (technical). Stars denote a one-sided test on ΔAUC (treatment vs matched control): * p < 0.05, ** p < 0.01, *** p < 0.002; NS otherwise. Primary inference in the main text uses AUC Days 2–7 (see Figure 2 and Figure S7).

**Figure S9:** Heatmap of transcriptome-wide expression. Columns are the 16 libraries grouped by treatment (Sulf, Ctrl, Curc, Comb; colored chips above). Values are row-wise z-scores of log2-FPKM. Rows and columns were clustered using Euclidean distance with ward.D2 linkage. The heatmap shows distinct global profiles for each treatment and coherent clustering of biological replicates.

**Figure S10:** Pairwise DEGs between treatments (*padj* ≤ 0.05; |log FC| ≥ 1). Bars show numbers of Up and Down DEGs for Curc–Sulf, Curc–Combo, and Sulf–Combo; totals are annotated above each pair.

**Figure S11:** Complete STRING protein-protein network of DEGs under Curcumin treatment. Edges: STRING v12.0, combined score ≥ 0.7; nodes are DEGs only (*padj* ≤ 0.05; |log2FC| ≥ 1); isolates removed; communities by Glay (node color = community; diamond node = Up-DEG; circle node = Down-DEG, node size = degree of the node).

**Figure S12:** Complete STRING protein-protein network of DEGs under Sulforaphane treatment. Edges: STRING v12.0, combined score ≥ 0.7; nodes are DEGs only (*padj* ≤ 0.05; |log2FC| ≥ 1); isolates removed; communities by Glay (node color = community; diamond node = Up-DEG; circle node = Down-DEG, node size = degree of the node).

**Figure S13:** Complete STRING protein-protein network of DEGs under Combination treatment. Edges: STRING v12.0, combined score ≥ 0.7; nodes are DEGs only (*padj* ≤ 0.05; |log2FC| ≥ 1); isolates removed; communities by Glay (node color = community; diamond node = Up-DEG; circle node = Down-DEG, node size = degree of the node).

### Supplementary Tables

**Table S1**: RNA-seq sample metadata and alignment metrics. For each Day–3 library (n = 4/condition; Control/Curcumin/Sulforaphane/Combination), the table reports total reads, alignment and annotation metrics. %mapped = total mapped / total reads; %unique = uniquely mapping reads; %multi = multi-mapping reads; %proper_pair = percent read pairs flagged by the aligner as properly mapped; %exon = reads mapped on to exon; %intron = reads mapped on to intron; %intergenic = reads mapped on to intergenic regions; here we report percent values (rounded to two decimals) and list total reads as counts.

**Table S2**: Differentially expressed genes: Curcumin vs control. DEGs for Curcumin relative to control (padj ≤ 0.05, |log FC| ≥ 1). Table provides the Entrez gene ID, gene name, log2FC, p-value, adjusted p-value (padj), and direction (Up/Down). The rows are sorted by padj (ascending) and then by |log2FC| (descending).

**Table S3**: Differentially expressed genes: Sulforaphane vs control. DEGs for Sulforaphane relative to control (padj ≤ 0.05, |log FC| ≥ 1). Table provides the Entrez gene ID, gene name, log2FC, p-value, adjusted p-value (padj), and direction (Up/Down). The rows are sorted by padj (ascending) and then by |log2FC| (descending).

**Table S4**: Differentially expressed genes: Combination vs control. DEGs for Combination relative to control (padj ≤ 0.05, |log FC| ≥ 1). Table provides the Entrez gene ID, gene name, log2FC, p-value, adjusted p-value (padj), and direction (Up/Down). The rows are sorted by padj (ascending) and then by |log2FC| (descending).

**Table S5**: Curcumin–Sulforaphane co-regulated and direction-reversed genes (vs control). Genes meeting the thresholds (padj ≤ 0.05, |log2FC| ≥ 1) in both contrasts are grouped as Co-up-regulated (n = 17) and Co-down-regulated (n = 16). Genes with opposite regulation between the contrasts are grouped as Curc (up) / Sulf (down) (n = 10) and Curc (down) / Sulf (up) (n = 5). Entries are C. elegans gene symbols; per-gene log2FC and padj for each contrast are provided in the DEG tables (Supplementary Tables S2–S4).

**Table S6**: Benchmarking overlap of Sulforaphane-responsive genes with Sedore et al. (2025). Up- and down-regulated DEGs for Sulforaphane vs control (SS104 glp-4(bn2), 0.625 mg/mL, Day 3 adults; padj ≤ 0.05; |log2FC| ≥ 1) compared with Sedore et al. (N2,100 µM sulforaphane) at Day 4 and Day 8. Table provides the overlapping Entrez gene IDs between Sedore set (Day 4 Up; Day 8 Up; Day 8 Down), and our Up/Down set. Summary: Day 4 Up ∩ our Up = 11/29 (38%); Day 4 Up ∩ our Down = 0/29 (0%); Day 8 Up ∩ our Up = 45/369 (12%); Day 8 Up ∩ our Down = 0/369 (0%); Day 8 Down ∩ our Down = 16/88 (18%); Day 8 Down ∩ our Up = 0/88 (0%). No direction mismatches were observed.

**Table S7**: Combination-unique Up-DEGs (padj ≤ 0.05; log2FC ≥ 1). Genes that are up-regulated in the Combination vs control and not differentially expressed in Curcumin vs control or Sulforaphane vs control at the same thresholds. The table provides Entrez gene id, gene name, log2FC (Combination vs control), and padj. Rows are sorted by padj (ascending) and then by |log2FC| (descending).

**Table S8**: Combination-unique Down-DEGs (padj ≤ 0.05; log2FC ≤ -1). Genes that are down-regulated in the Combination vs control and not differentially expressed in Curcumin vs control or Sulforaphane vs control at the same thresholds. The table provides Entrez gene id, gene name, log2FC (Combination vs control), and padj. Rows are sorted by padj (ascending) and then by |log2FC| (descending).

**Table S9**: GO Biological Process enrichment (clusterProfiler). GO BP enrichment for Curc, Sulf, and Comb versus control was performed with clusterProfiler. Terms were retained at padj ≤ 0.05 and qvalue ≤ 0.05; analyses used a common background (union of expressed genes) and UP/DOWN gene sets were analyzed separately. Columns report the Direction (UP/DOWN), Set (Curc/Sulf/Comb vs Ctrl), GO ID, Description, hit Count, GeneRatio (Count/input list size), BgRatio (term/background size), pvalue, p.adjust, qvalue and the geneID list (slash-separated gene names).

**Table S10**: KEGG pathway enrichment (clusterProfiler). KEGG pathway enrichment for Curc, Sulf, and Comb versus control was performed with clusterProfiler. Terms were retained at padj ≤ 0.05 and qvalue ≤ 0.05; analyses used a common background (union of expressed genes) and UP/DOWN gene sets were analyzed separately. Columns report the Direction (UP/DOWN), Set (Curc/Sulf/Comb vs Ctrl), Pathway ID, Description, hit Count, GeneRatio (Count/input list size), BgRatio (term/background size), pvalue, p.adjust, qvalue and the geneID list (slash-separated Entrez gene IDs).

**Table S11**: CelEst transcription-factor activity inference per treatment. Blocks list TF activity for Curc, Sulf, and Combo versus control. Columns report the TF gene symbol, Entrez gene ID, CelEst activity score (score; positive = activation, negative = repression), p-value, and adjusted p-value (padj). Only TFs with padj < 0.1 are shown; the full CelEst output is available on request.

**Table S12**: Node attributes for STRING protein-protein interaction network (by treatment). Nodes correspond to DEGs (padj ≤ 0.05; |log2FC| ≥ 1) used in the STRING v12.0 association networks (combined score ≥ 0.7; DEGs-only; isolates removed; GLay communities). Columns report gene symbol, Entrez gene ID, log2FC (vs control), direction (Up/Down), degree, GLay cluster ID, and in_LCC (membership in the largest connected component). One sheet includes Curc, Sulf, and Combo (separated by blocks: Curc vs Ctrl, Sulf vs Ctrl, Comb vs Ctrl).

**Table S13**: Selected genes and module assignment for the curated 50-gene heatmap. For each gene used in Figure 6, the table lists the gene symbol, Entrez ID, module (Glutathione/Detox; Sphingolipid/Lipid; Innate immunity; Longevity/Stress), and the selection rank based on maximum |log2FC| across treatments (Curc, Sulf, Comb vs Ctrl). Genes appearing in multiple candidate blocks were assigned to the block in which they achieved the highest rank. Module definitions (GO/KEGG):Glutathione/Detox: KEGG cel00480 (Glutathione metabolism), cel00980 (Metabolism of xenobiotics by cytochrome P450), cel00982 (Drug metabolism cytochrome P450); GO GO:0006749 (glutathione metabolic process), GO:0006805 (xenobiotic metabolic process), GO:0098869 (cellular oxidant detoxification). Sphingolipid/Lipid: KEGG cel00600 (Sphingolipid metabolism), cel01040 (Biosynthesis of unsaturated fatty acids); GO GO:0006665 (sphingolipid metabolic process), GO:0030148 (sphingolipid biosynthetic process), GO:0046466 (membrane lipid catabolic process). Innate immunity: GO GO:0009607 (response to biotic stimulus), GO:0098542 (defense response to other organism), GO:0045087 (innate immune response). Longevity/Stress: KEGG cel04212 (Longevity regulating pathway worm), cel04142 (Lysosome); GO GO:0098771 (inorganic ion homeostasis).

## References

Alves, Inês, Edilene Maria Queiroz Araújo, Louise T. Dalgaard, Sharda Singh, Elisabet Børsheim, and Eugenia Carvalho. 2025. “Protective Effects of Sulforaphane Preventing Inflammation and Oxidative Stress to Enhance Metabolic Health: A Narrative Review.” Nutrients 17 (3): 428. 10.3390/nu17030428.

An, Jae Hyung, and T. Keith Blackwell. 2003. “SKN-1 Links *C. Elegans* Mesendodermal Specification to a Conserved Oxidative Stress Response.” Genes & Development 17 (15): 1882–93. 10.1101/gad.1107803.

Argyropoulou, Aikaterini, Nektarios Aligiannis, Ioannis P. Trougakos, and Alexios-Leandros Skaltsounis. 2013. “Natural Compounds with Anti-Ageing Activity.” Natural Product Reports 30 (11): 1412. 10.1039/c3np70031c.

Ashburner, Michael, Catherine A. Ball, Judith A. Blake, et al. 2000. “Gene Ontology: Tool for the Unification of Biology.” Nature Genetics 25 (1): 25–29. 10.1038/75556.

Bjørklund, Geir, Mariia Shanaida, Roman Lysiuk, et al. 2022. “Natural Compounds and Products from an Anti-Aging Perspective.” Molecules 27 (20): 7084. 10.3390/molecules27207084.

Blackwell, T. Keith, Michael J. Steinbaugh, John M. Hourihan, Collin Y. Ewald, and Meltem Isik. 2015. “SKN-1/Nrf, Stress Responses, and Aging in *Caenorhabditis Elegans*.” Free Radical Biology and Medicine, Nrf2 Regulated Redox Signaling and Metabolism in Physiology and Medicine, vol. 88 (November): 290–301. 10.1016/j.freeradbiomed.2015.06.008.

Cheng, A. L., C. H. Hsu, J. K. Lin, et al. 2001. “Phase I Clinical Trial of Curcumin, a Chemopreventive Agent, in Patients with High-Risk or Pre-Malignant Lesions.” Anticancer Research 21 (4B): 2895–900.

Cheng, Maojun, Fang Ding, Liyang Li, et al. 2025. “Exploring the Role of Curcumin in Mitigating Oxidative Stress to Alleviate Lipid Metabolism Disorders.” Frontiers in Pharmacology 16 (January). 10.3389/fphar.2025.1517174.

Churgin, Matthew A., Sang-Kyu Jung, Chih-Chieh Yu, Xiangmei Chen, David M. Raizen, and Christopher Fang-Yen. 2017. “Longitudinal Imaging of Caenorhabditis Elegans in a Microfabricated Device Reveals Variation in Behavioral Decline during Aging.” eLife 6 (May): e26652. 10.7554/eLife.26652.

Clarke, John D., Anna Hsu, David E. Williams, et al. 2011. “Metabolism and Tissue Distribution of Sulforaphane in Nrf2 Knockout and Wild-Type Mice.” Pharmaceutical Research 28 (12): 3171–79. 10.1007/s11095-011-0500-z.

Collins, James J., Cheng Huang, Stacie Hughes, and Kerry Kornfeld. 2018. “The Measurement and Analysis of Age-Related Changes in Caenorhabditis Elegans.” In WormBook: The Online Review of C. Elegans Biology [Internet]. WormBook. https://www.ncbi.nlm.nih.gov/books/NBK116075/.

Conway, Jake R., Alexander Lex, and Nils Gehlenborg. 2017. “UpSetR: An R Package for the Visualization of Intersecting Sets and Their Properties.” Bioinformatics 33 (18): 2938–40. 10.1093/bioinformatics/btx364.

Cutler, Roy G., Kenneth W. Thompson, Simonetta Camandola, Kendra T. Mack, and Mark P. Mattson. 2014. “Sphingolipid Metabolism Regulates Development and Lifespan in Caenorhabditis Elegans.” Mechanisms of Ageing and Development 143–144 (December): 9–18. 10.1016/j.mad.2014.11.002.

Davinelli, Sergio, Alessandro Medoro, Frank B. Hu, and Giovanni Scapagnini. 2025. “Dietary Polyphenols as Geroprotective Compounds: From Blue Zones to Hallmarks of Ageing.” Ageing Research Reviews 108 (June): 102733. 10.1016/j.arr.2025.102733.

Dinkova-Kostova, Albena T., Jed W. Fahey, Rumen V. Kostov, and Thomas W. Kensler. 2017. “KEAP1 and Done? Targeting the NRF2 Pathway with Sulforaphane.” Trends in Food Science & Technology 69 (November): 257–69. 10.1016/j.tifs.2017.02.002.

Edwards, Clare, John Canfield, Neil Copes, Muhammad Rehan, David Lipps, and Patrick C. Bradshaw. 2014. “D-Beta-Hydroxybutyrate Extends Lifespan in C. Elegans.” Aging 6 (8): 621–44. 10.18632/aging.100683.

Egner, Patricia A., Jian Guo Chen, Jin Bing Wang, et al. 2011. “Bioavailability of Sulforaphane from Two Broccoli Sprout Beverages: Results of a Short-Term, Cross-over Clinical Trial in Qidong, China.” Cancer Prevention Research 4 (3): 384–95. 10.1158/1940-6207.CAPR-10-0296.

Egner, Patricia A., Jian-Guo Chen, Adam T. Zarth, et al. 2014. “Rapid and Sustainable Detoxication of Airborne Pollutants by Broccoli Sprout Beverage: Results of a Randomized Clinical Trial in China.” Cancer Prevention Research 7 (8): 813–23. 10.1158/1940-6207.CAPR-14-0103.

Fang, Evandro F., Tyler B. Waltz, Henok Kassahun, et al. 2017. “Tomatidine Enhances Lifespan and Healthspan in C. Elegans through Mitophagy Induction via the SKN-1/Nrf2 Pathway.” Scientific Reports 7 (1): 46208. 10.1038/srep46208.

Franceschi, C., and J. Campisi. 2014. “Chronic Inflammation (Inflammaging) and Its Potential Contribution to Age-Associated Diseases.” The Journals of Gerontology Series A: Biological Sciences and Medical Sciences 69 (Suppl 1): S4–9. 10.1093/gerona/glu057.

Gaesser, Glenn A., Stephanie E. Hall, Siddhartha S. Angadi, David C. Poole, and Susan B. Racette. 2025. “Increasing the Health Span: Unique Role for Exercise.” Journal of Applied Physiology 138 (6): 1285–308. 10.1152/japplphysiol.00049.2025.

Glenn, Charles F., David K. Chow, Lawrence David, et al. 2004. “Behavioral Deficits During Early Stages of Aging in Caenorhabditis Elegans Result From Locomotory Deficits Possibly Linked to Muscle Frailty.” The Journals of Gerontology: Series A 59 (12): 1251–60. 10.1093/gerona/59.12.1251.

Guralnik, J. M., E. M. Simonsick, L. Ferrucci, et al. 1994. “A Short Physical Performance Battery Assessing Lower Extremity Function: Association with Self-Reported Disability and Prediction of Mortality and Nursing Home Admission.” Journal of Gerontology 49 (2): M85–94. 10.1093/geronj/49.2.m85.

Guralnik, Jack M., Luigi Ferrucci, Carl F. Pieper, et al. 2000. “Lower Extremity Function and Subsequent Disability: Consistency Across Studies, Predictive Models, and Value of Gait Speed Alone Compared With the Short Physical Performance Battery.” The Journals of Gerontology: Series A 55 (4): M221–31. 10.1093/gerona/55.4.M221.

Hahm, Jeong-Hoon, Sunhee Kim, Race DiLoreto, et al. 2015. “C. Elegans Maximum Velocity Correlates with Healthspan and Is Maintained in Worms with an Insulin Receptor Mutation.” Nature Communications 6 (1): 8919. 10.1038/ncomms9919.

Herndon, Laura A., Peter J. Schmeissner, Justyna M. Dudaronek, et al. 2002. “Stochastic and Genetic Factors Influence Tissue-Specific Decline in Ageing C. Elegans.” Nature 419 (6909): 808–14. 10.1038/nature01135.

Huang, Cheng, Chengjie Xiong, and Kerry Kornfeld. 2004. “Measurements of Age-Related Changes of Physiological Processes That Predict Lifespan of Caenorhabditis Elegans.” Proceedings of the National Academy of Sciences 101 (21): 8084–89. 10.1073/pnas.0400848101.

Ji, Huihui, Zhimin Qi, Daniel Schrapel, et al. 2021. “Sulforaphane Targets TRA-1/GLI Upstream of DAF-16/FOXO to Promote C. Elegans Longevity and Healthspan.” Frontiers in Cell and Developmental Biology 9 (December): 784999. 10.3389/fcell.2021.784999.

Jones, Owen R., Alexander Scheuerlein, Roberto Salguero-Gómez, et al. 2014. “Diversity of Ageing across the Tree of Life.” Nature 505 (7482): 169–73. 10.1038/nature12789.

Kaletta, Titus, and Michael O. Hengartner. 2006. “Finding Function in Novel Targets: C. Elegans as a Model Organism.” Nature Reviews Drug Discovery 5 (5): 387–99. 10.1038/nrd2031.

Kanehisa, Minoru, Miho Furumichi, Yoko Sato, Yuriko Matsuura, and Mari Ishiguro-Watanabe. 2025. “KEGG: Biological Systems Database as a Model of the Real World.” Nucleic Acids Research 53 (D1): D672–77. 10.1093/nar/gkae909.

Karimian, Maryam S., Matteo Pirro, Thomas P. Johnston, Muhammed Majeed, and Amirhossein Sahebkar. 2017. “Curcumin and Endothelial Function: Evidence and Mechanisms of Protective Effects.” Current Pharmaceutical Design 23 (17). 10.2174/1381612823666170222122822.

Kawamura, Kazuto, and Ichiro N. Maruyama. 2019. “Forward Genetic Screen for *Caenorhabditis Elegans* Mutants with a Shortened Locomotor Healthspan.” G3 Genes|Genomes|Genetics 9 (8): 2415–23. 10.1534/g3.119.400241.

Kennedy, Brian K., Shelley L. Berger, Anne Brunet, et al. 2014. “Geroscience: Linking Aging to Chronic Disease.” Cell 159 (4): 709–13. 10.1016/j.cell.2014.10.039.

Kenyon, Cynthia J. 2010. “The Genetics of Ageing.” Nature 464 (7288): 504–12. 10.1038/nature08980.

Kim, Daehwan, Joseph M. Paggi, Chanhee Park, Christopher Bennett, and Steven L. Salzberg. 2019. “Graph-Based Genome Alignment and Genotyping with HISAT2 and HISAT-Genotype.” Nature Biotechnology 37 (8): 907–15. 10.1038/s41587-019-0201-4.

Kim, Yong Sook, Youngkeun Ahn, Moon Hwa Hong, et al. 2007. “Curcumin Attenuates Inflammatory Responses of TNF-α-Stimulated Human Endothelial Cells.” Journal of Cardiovascular Pharmacology 50 (1): 41–49. 10.1097/FJC.0b013e31805559b9.

Kirchweger, Benjamin, Julia Zwirchmayr, Ulrike Grienke, and Judith M. Rollinger. 2023. “The Role of Caenorhabditis Elegans in the Discovery of Natural Products for Healthy Aging.” Natural Product Reports 40 (12): 1849–73. 10.1039/D3NP00021D.

Kirkwood, Thomas B. L., and Steven N. Austad. 2000. “Why Do We Age?” Nature 408 (6809): 233–38. 10.1038/35041682.

Kolde, Raivo. 2025. *Pheatmap: Pretty Heatmap*s. https://github.com/raivokolde/pheatmap.

Lao, Christopher D., Mack T. Ruffin, Daniel Normolle, et al. 2006. “Dose Escalation of a Curcuminoid Formulation.” BMC Complementary and Alternative Medicine 6 (1): 10. 10.1186/1472-6882-6-10.

Lapierre, Louis R., Sara Gelino, Alicia Meléndez, and Malene Hansen. 2011. “Autophagy and Lipid Metabolism Coordinately Modulate Life Span in Germline-Less C. Elegans.” Current Biology 21 (18): 1507–14. 10.1016/j.cub.2011.07.042.

Lee, Da-Yeon, Su-Jeong Lee, Prabha Chandrasekaran, et al. 2023. “Dietary Curcumin Attenuates Hepatic Cellular Senescence by Suppressing the MAPK/NF-κB Signaling Pathway in Aged Mice.” Antioxidants 12 (6): 1165. 10.3390/antiox12061165.

Lex, Alexander, Nils Gehlenborg, Hendrik Strobelt, Romain Vuillemot, and Hanspeter Pfister. 2014. “UpSet: Visualization of Intersecting Sets.” IEEE Transactions on Visualization and Computer Graphics 20 (12): 1983–92. 10.1109/TVCG.2014.2346248.

Liao, Yang, Gordon K. Smyth, and Wei Shi. 2014. “featureCounts: An Efficient General Purpose Program for Assigning Sequence Reads to Genomic Features.” Bioinformatics 30 (7): 923–30. 10.1093/bioinformatics/btt656.

Lin, Yugui, Chunxiu Lin, Yong Cao, and Yunjiao Chen. 2023. “Caenorhabditis Elegans as an in Vivo Model for the Identification of Natural Antioxidants with Anti-Aging Actions.” Biomedicine & Pharmacotherapy 167 (November): 115594. 10.1016/j.biopha.2023.115594.

Liu, Shuyan, Nadine Saul, Bo Pan, Ralph Menzel, and Christian E. W. Steinberg. 2013. “The Non-Target Organism Caenorhabditis Elegans Withstands the Impact of Sulfamethoxazole.” Chemosphere 93 (10): 2373–80. 10.1016/j.chemosphere.2013.08.036.

López-Otín, Carlos, Maria A. Blasco, Linda Partridge, Manuel Serrano, and Guido Kroemer. 2013. “The Hallmarks of Aging.” Cell 153 (6): 1194–217. 10.1016/j.cell.2013.05.039.

Love, Michael I., Wolfgang Huber, and Simon Anders. 2014. “Moderated Estimation of Fold Change and Dispersion for RNA-Seq Data with DESeq2.” Genome Biology 15 (12): 550. 10.1186/s13059-014-0550-8.

Mathew, Mark D., Neal D. Mathew, and Paul R. Ebert. 2012. “WormScan: A Technique for High-Throughput Phenotypic Analysis of Caenorhabditis Elegans.” PLOS ONE 7 (3): e33483. 10.1371/journal.pone.0033483.

Mohanraj, Karthikeyan, Bagavathy Shanmugam Karthikeyan, R. P. Vivek-Ananth, et al. 2018. “IMPPAT: A Curated Database of Indian Medicinal Plants, Phytochemistry And Therapeutics.” Scientific Reports 8 (1): 4329. 10.1038/s41598-018-22631-z.

Newell Stamper, Breanne L., James R. Cypser, Katerina Kechris, David Alan Kitzenberg, Patricia M. Tedesco, and Thomas E. Johnson. 2018. “Movement Decline across Lifespan of *Caenorhabditis Elegans* Mutants in the Insulin/Insulin-like Signaling Pathway.” Aging Cell 17 (1): e12704. 10.1111/acel.12704.

Newman, John C., and Eric Verdin. 2017. “β-Hydroxybutyrate: A Signaling Metabolite.” Annual Review of Nutrition 37 (1): 51–76. 10.1146/annurev-nutr-071816-064916.

Niccoli, Teresa, and Linda Partridge. 2012. “Ageing as a Risk Factor for Disease.” Current Biology 22 (17): R741–52. 10.1016/j.cub.2012.07.024.

Ogawa, Takahiro, Yukihiro Kodera, Dai Hirata, T. Keith Blackwell, and Masaki Mizunuma. 2016. “Natural Thioallyl Compounds Increase Oxidative Stress Resistance and Lifespan in Caenorhabditis Elegans by Modulating SKN-1/Nrf.” Scientific Reports 6 (1): 21611. 10.1038/srep21611.

O’Reilly, Linda P., Cliff J. Luke, David H. Perlmutter, Gary A. Silverman, and Stephen C. Pak. 2014. “*C. Elegans* in High-Throughput Drug Discovery.” *Advanced Drug Delivery Reviews*, Innovative tissue models for drug discovery and development, vols. 69–70 (April): 247–53. 10.1016/j.addr.2013.12.001.

Perez, Marcos Francisco. 2024. Coding-Free Differential TF Activity Estimation from C. Elegans Transcriptomic Data with the CelEsT R Shiny*…* November 21. https://www.protocols.io/view/coding-free-differential-tf-activity-estimation-fr-dmxq47mw.

Perez, Marcos Francisco. 2025. “CelEst: A Unified Gene Regulatory Network for Estimating Transcription Factor Activities in C. Elegans.” Genetics 229 (3): iyae189. 10.1093/genetics/iyae189.

Pertea, Mihaela, Geo M. Pertea, Corina M. Antonescu, Tsung-Cheng Chang, Joshua T. Mendell, and Steven L. Salzberg. 2015. “StringTie Enables Improved Reconstruction of a Transcriptome from RNA-Seq Reads.” Nature Biotechnology 33 (3): 290–95. 10.1038/nbt.3122.

Rantakokko, Merja, Minna Mänty, and Taina Rantanen. 2013. “Mobility Decline in Old Age.” Exercise and Sport Sciences Reviews 41 (1): 19. 10.1097/JES.0b013e3182556f1e.

Sedore, Christine A., Erik Segerdell, Anna L. Coleman-Hulbert, et al. 2025. “The Broccoli Derivative Sulforaphane Extends Lifespan by Slowing the Transcriptional Aging Clock.” Preprint, bioRxiv, May 15. 10.1101/2025.05.11.653363.

Shannon, Paul, Andrew Markiel, Owen Ozier, et al. 2003. “Cytoscape: A Software Environment for Integrated Models of Biomolecular Interaction Networks.” Genome Research 13 (11): 2498–504. 10.1101/gr.1239303.

Shen, Li-Rong, Laurence D. Parnell, Jose M. Ordovas, and Chao-Qiang Lai. 2013. “Curcumin and Aging.” BioFactors 39 (1): 133–40. 10.1002/biof.1086.

Shimazu, Tadahiro, Matthew D. Hirschey, John Newman, et al. 2013. “Suppression of Oxidative Stress by β-Hydroxybutyrate, an Endogenous Histone Deacetylase Inhibitor.” Science 339 (6116): 211–14. 10.1126/science.1227166.

Studenski, Stephanie, Subashan Perera, Kushang Patel, et al. 2011. “Gait Speed and Survival in Older Adults.” JAMA 305 (1): 50–58. 10.1001/jama.2010.1923.

Suzuki, Takafumi, Jun Takahashi, and Masayuki Yamamoto. 2023. “Molecular Basis of the KEAP1-NRF2 Signaling Pathway.” Molecules and Cells 46 (3): 133–41. 10.14348/molcells.2023.0028.

Szklarczyk, Damian, Rebecca Kirsch, Mikaela Koutrouli, et al. 2023. “The STRING Database in 2023: Protein–Protein Association Networks and Functional Enrichment Analyses for Any Sequenced Genome of Interest.” Nucleic Acids Research 51 (D1): D638–46. 10.1093/nar/gkac1000.

The Gene Ontology Consortium, Suzi A. Aleksander, James Balhoff, et al. 2023. “The Gene Ontology Knowledgebase in 2023.” Genetics 224 (1): iyad031. 10.1093/genetics/iyad031.

Tissenbaum, Heidi A. 2012. “Genetics, Life Span, Health Span, and the Aging Process in Caenorhabditis Elegans.” The Journals of Gerontology: Series A 67A (5): 503–10. 10.1093/gerona/gls088.

Virk, Bhupinder, Gonçalo Correia, David P. Dixon, et al. 2012. “Excessive Folate Synthesis Limits Lifespan in the C. Elegans: E. Coliaging Model.” BMC Biology 10 (1): 67. 10.1186/1741-7007-10-67.

Vivek-Ananth, R. P., Karthikeyan Mohanraj, Ajaya Kumar Sahoo, and Areejit Samal. 2023. “IMPPAT 2.0: An Enhanced and Expanded Phytochemical Atlas of Indian Medicinal Plants.” ACS Omega 8 (9): 8827–45. 10.1021/acsomega.3c00156.

Vural, Dervis C., Greg Morrison, and L. Mahadevan. 2014. “Aging in Complex Interdependency Networks.” Physical Review E 89 (2): 022811. 10.1103/PhysRevE.89.022811.

Weinkove, David, and Giulia Zavagno. 2021. “Applying C. Elegans to the Industrial Drug Discovery Process to Slow Aging.” Frontiers in Aging 2 (October): 740582. 10.3389/fragi.2021.740582.

Wink, Michael. 2022. “Current Understanding of Modes of Action of Multicomponent Bioactive Phytochemicals: Potential for Nutraceuticals and Antimicrobials.” Annual Review of Food Science and Technology 13 (1): 337–59. 10.1146/annurev-food-052720-100326.

World Health Organization. 2015. World Report on Ageing and Health. World Health Organization. https://iris.who.int/handle/10665/186463.

Wu, Tianzhi, Erqiang Hu, Shuangbin Xu, et al. 2021. “clusterProfiler 4.0: A Universal Enrichment Tool for Interpreting Omics Data.” The Innovation 2 (3). 10.1016/j.xinn.2021.100141.

Xu, Jianing, Pengyun Du, Xiaoyu Liu, Xiao Xu, Yuting Ge, and Chenggang Zhang. 2023. “Curcumin Supplementation Increases Longevity and Antioxidant Capacity in Caenorhabditis Elegans.” Frontiers in Pharmacology 14 (June): 1195490. 10.3389/fphar.2023.1195490.

Xu, Yiming, and Ling Liu. 2017. “Curcumin Alleviates Macrophage Activation and Lung Inflammation Induced by Influenza Virus Infection through Inhibiting the NF -κB Signaling Pathway.” Influenza and Other Respiratory Viruses 11 (5): 457–63. 10.1111/irv.12459.

Zavagno, Giulia, Adelaide Raimundo, Andy Kirby, Christopher Saunter, and David Weinkove. 2024. “Rapid Measurement of Ageing by Automated Monitoring of Movement of C. Elegans Populations.” GeroScience 46 (2): 2281–93. 10.1007/s11357-023-00998-w.

