## Supplementary Figures S1-S13 for "Curcumin and Sulforaphane Preserve Mobility in Aging *Caenorhabditis elegans* via Distinct yet Complementary Transcriptional Signatures"

**Curcumin and Sulforaphane Preserve Mobility in Aging *Caenorhabditis* elegans via Distinct yet Complementary Transcriptional Signatures**

R. P. Vivek-Ananth ^a,b^, Durai Sellegounder ^a^, Karthik Mohanraj ^c^, Sushmita Maitra ^c^,

Chris Saunter ^c^, David Weinkove ^c,d^, Eric Verdin ^a^, Stephen M. Phipps ^b^, Nathan D. Price ^a,b,*^

^a^ Buck Institute for Research on Aging, Novato, CA 94945, USA

^b^ Thorne HealthTech, New York, NY 10019, USA

^c^ Magnitude Biosciences Ltd., NETPark Plexus, Thomas Wright Way, Sedgefield TS21 3FD, UK

^d^ Department of Biosciences, Durham University, Stockton Road, Durham DH1 3LE, UK

* Authors to whom correspondence should be addressed:

Nathan D. Price

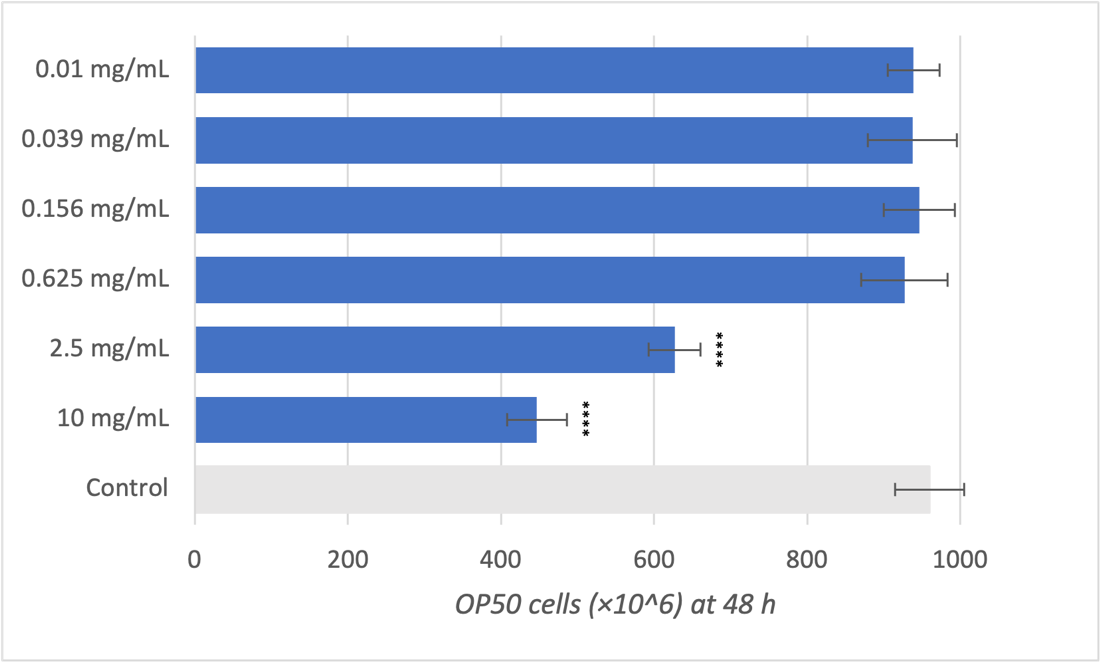

**Figure S1**: OP50 growth under sulforaphane for dose selection (48 h). Bars show mean ± SD cell counts (cells ×10⁶), n = 8 technical replicates per condition. One-way ANOVA with Dunnett’s adjusted comparisons vs water control: only 2.5 and 10 mg/mL sulforaphane significantly reduced OP50 counts (adjusted p < 0.0001); doses ≤ 0.625 mg/mL were not significant. Accordingly, 0.625 mg/mL was used as the OP50-compatible concentration in worm assays.

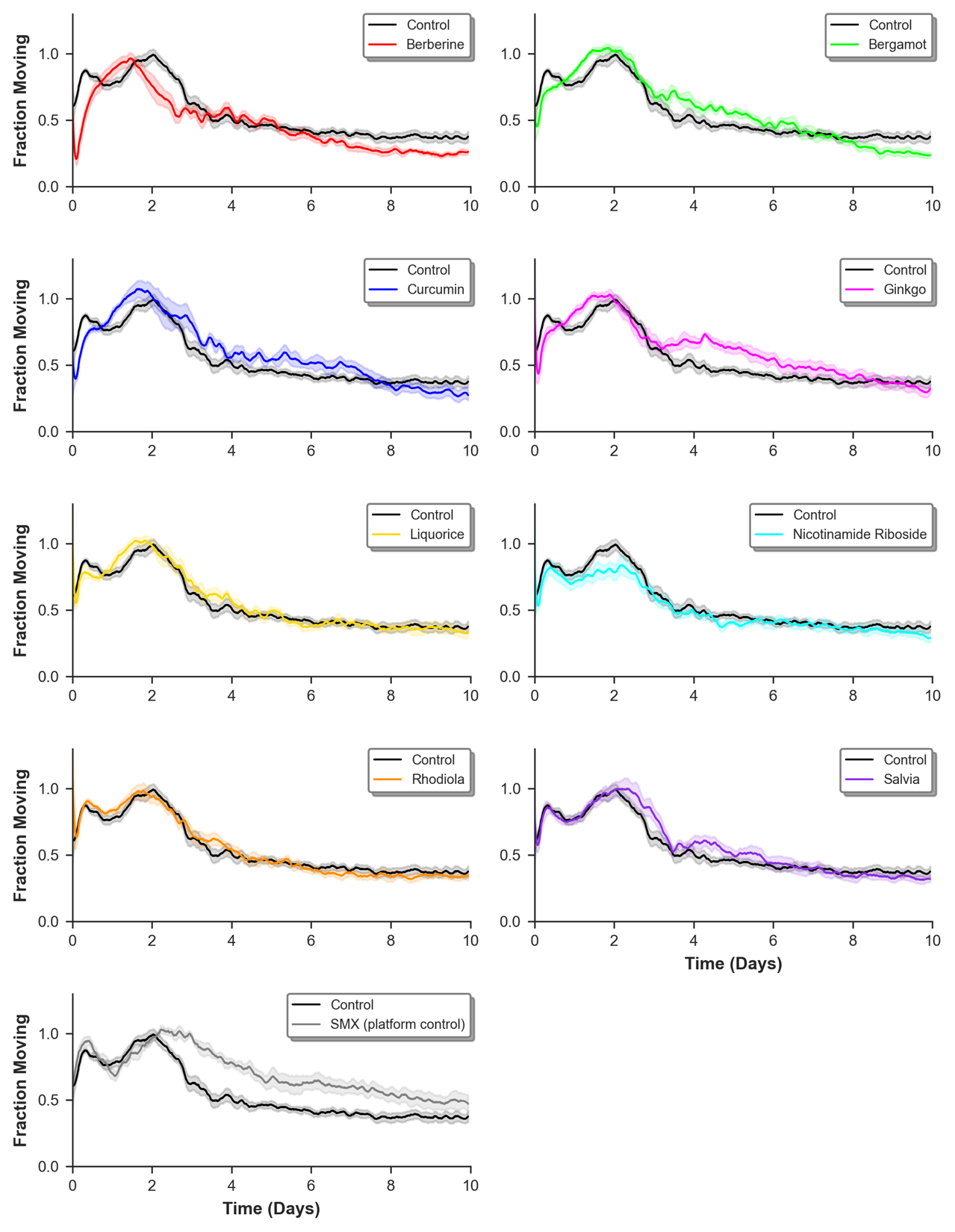

**Figure S2**: Fraction-moving trajectories for eight interventions and SMX (platform control), each overlaid with its matched control (gray). Curves span 0–10 days; AUC summaries over the post–Day 2 interval (48–238 h) are reported in the main text (Fig. 1c). SMX is shown to document assay sensitivity and is not used for head-to-head ranking.

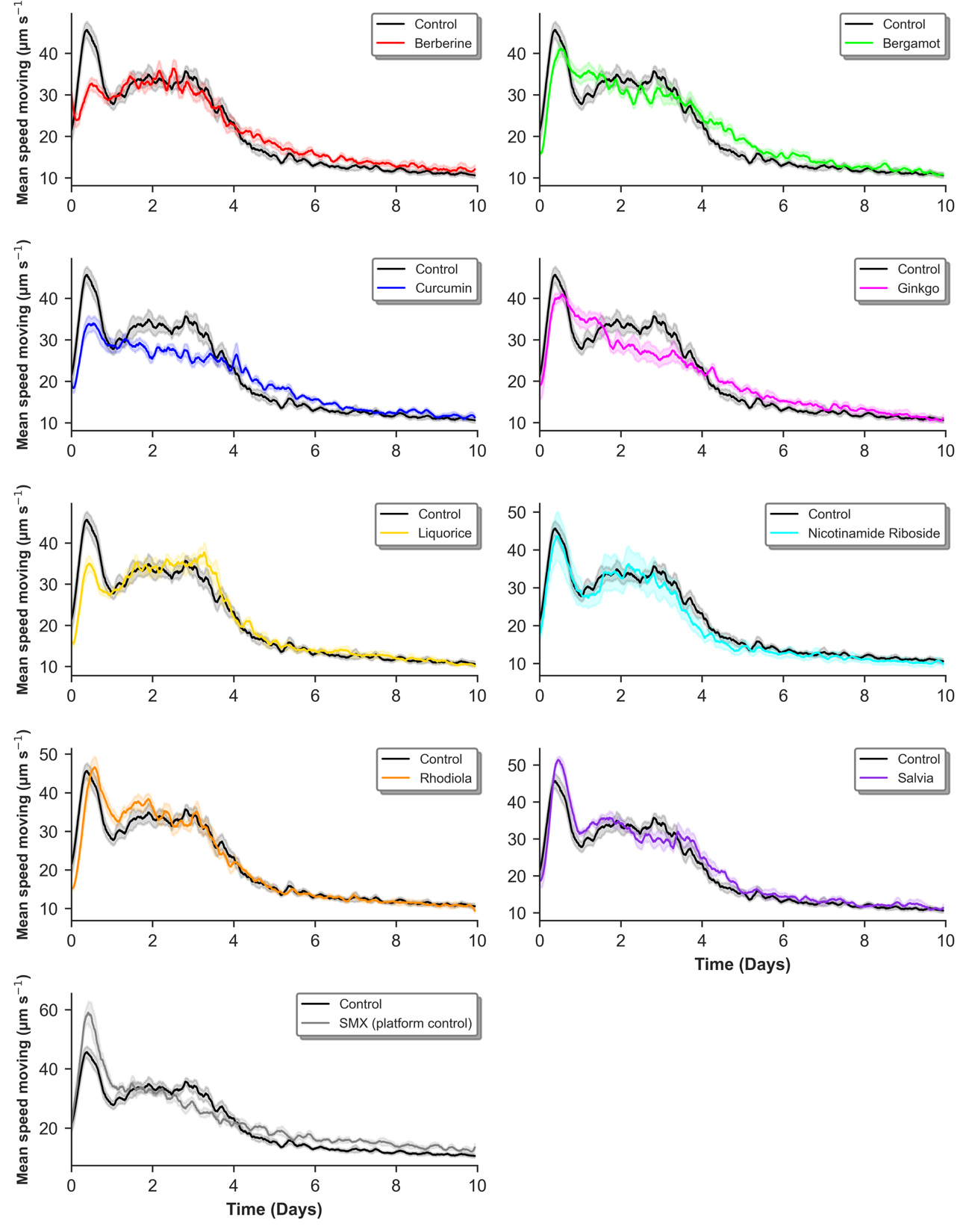

**Figure S3**: Speed of moving trajectories for eight interventions and SMX (platform control), each overlaid with its matched control (gray). Curves span 0–10 days; AUC summaries over the post–Day 2 interval (48–238 h) are reported in the main text (Fig. 1c). SMX is shown to document assay sensitivity and is not used for head-to-head ranking.

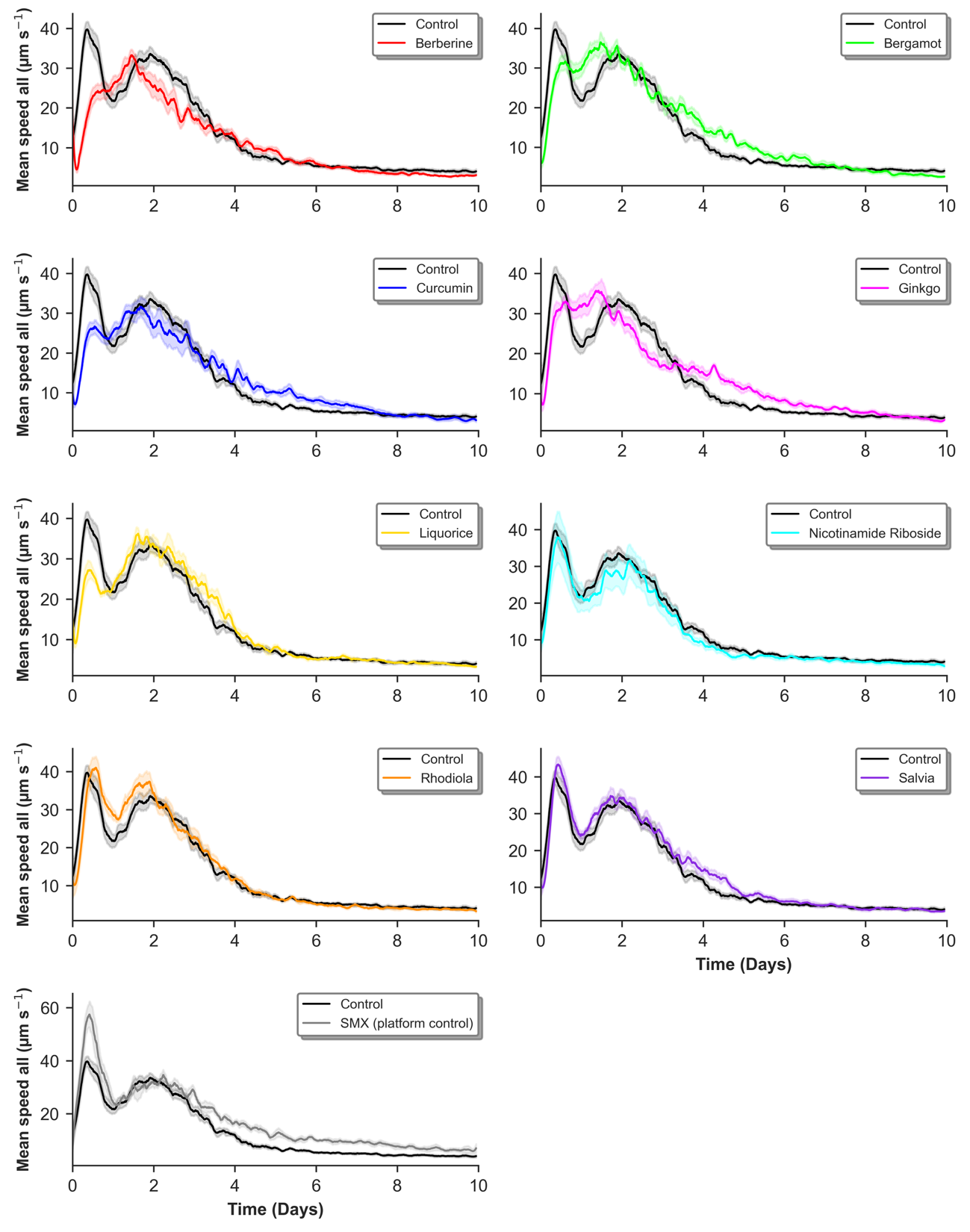

**Figure S4**: Speed of all worms trajectories for eight interventions and SMX (platform control), each overlaid with its matched control (gray). Curves span 0–10 days; AUC summaries over the post–Day 2 interval (48–238 h) are reported in the main text (Fig. 1c). SMX is shown to document assay sensitivity and is not used for head-to-head ranking.

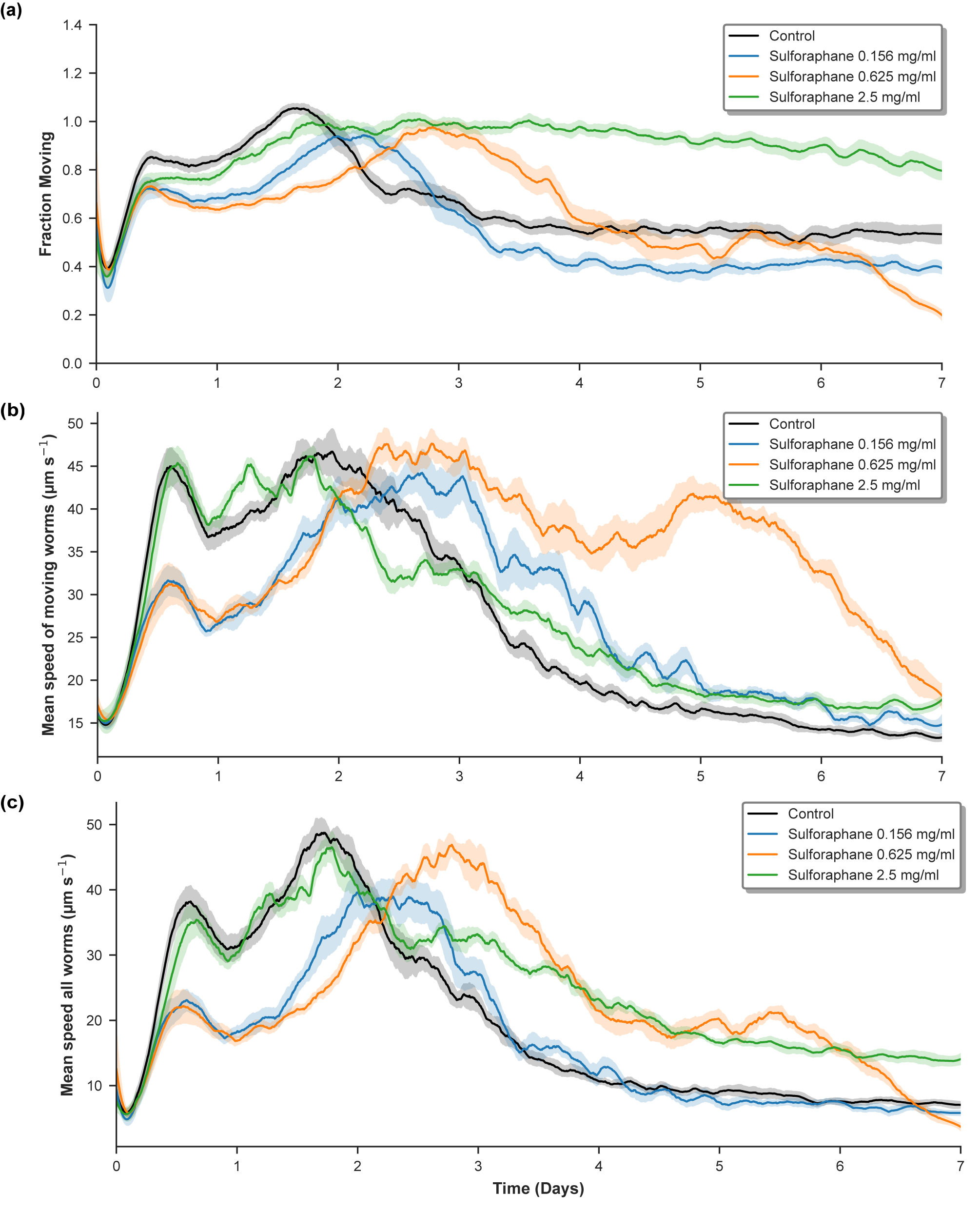

**Figure S5**: Sulforaphane dose comparison in the screen. (a) Fraction moving, (b) speed of moving (µm·s⁻¹), and (c) mean speed of all worms (µm·s⁻¹) for control and sulforaphane at 0.156, 0.625, and 2.5 mg/mL.

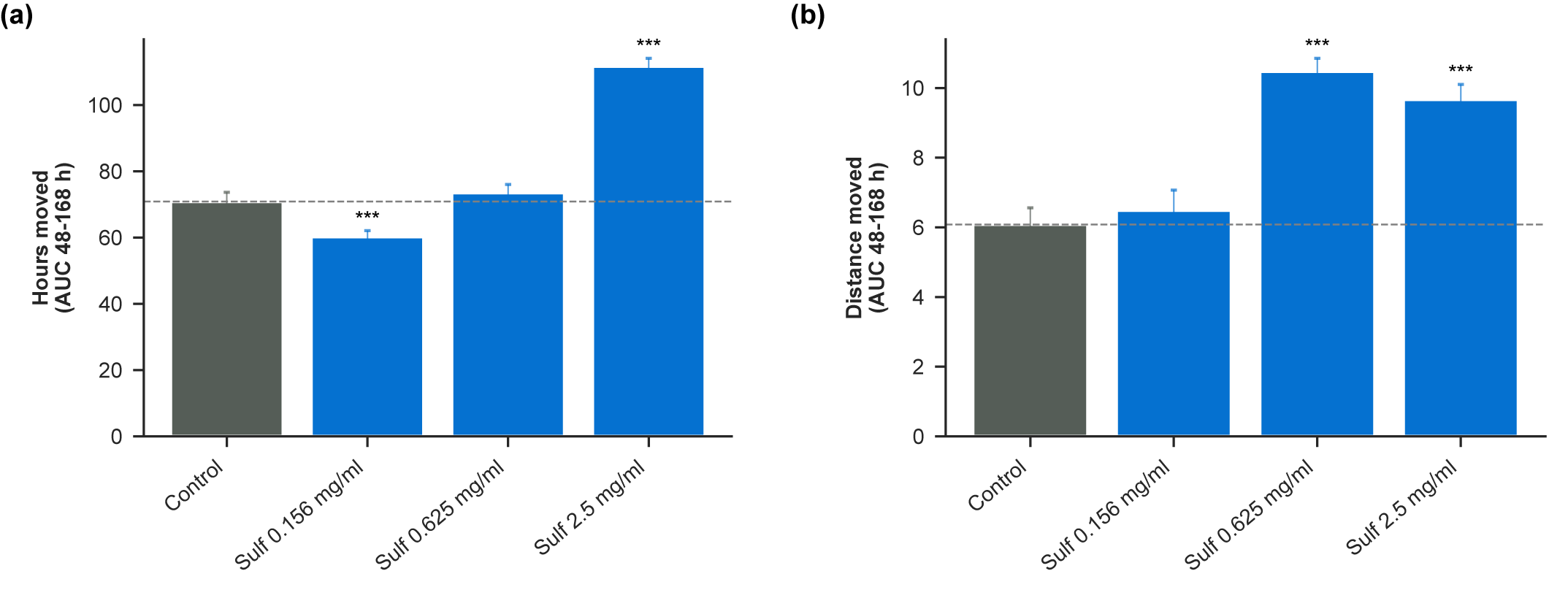

**Figure S6**: Sulforaphane dose comparison in the screen: AUC endpoints (48–168 hours). (a) Hours moved (AUC of fraction moving) and (b) distance moved (AUC of mean speed of all worms) for sulforaphane 0.156, 0.625, and 2.5 mg/mL versus the matched control (bars = mean ± SEM from technical replicates). Stars denote a one-sided test on ΔAUC (treatment vs matched control): * p < 0.05, ** p < 0.01, *** p < 0.002; NS otherwise.

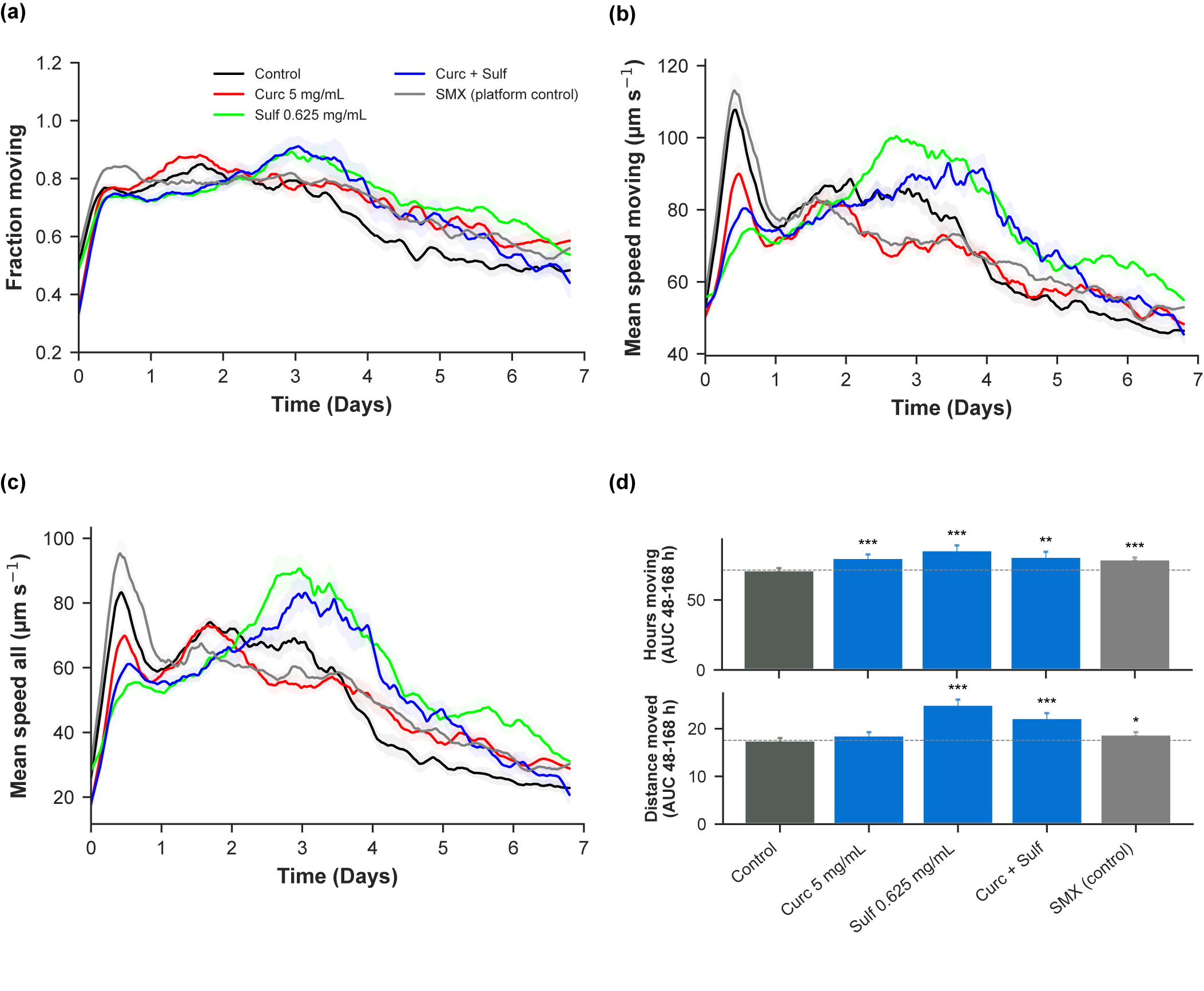

**Figure S7**: Mobility under curcumin (5 mg/mL), sulforaphane (0.625 mg/mL), and their combination. (a–c) Time courses for fraction moving, mean speed of moving, and mean speed of all worms (Replicate 2; mean ± SEM shown). SMX is a platform control. (d) AUC (48–168 h) for hours moved and distance moved relative to matched controls. Stars denote a one-sided test on ΔAUC (treatment vs matched control): * p < 0.05, ** p < 0.01, *** p < 0.002; NS otherwise. Replicate 1 is shown in Figure 2 (identical layout).

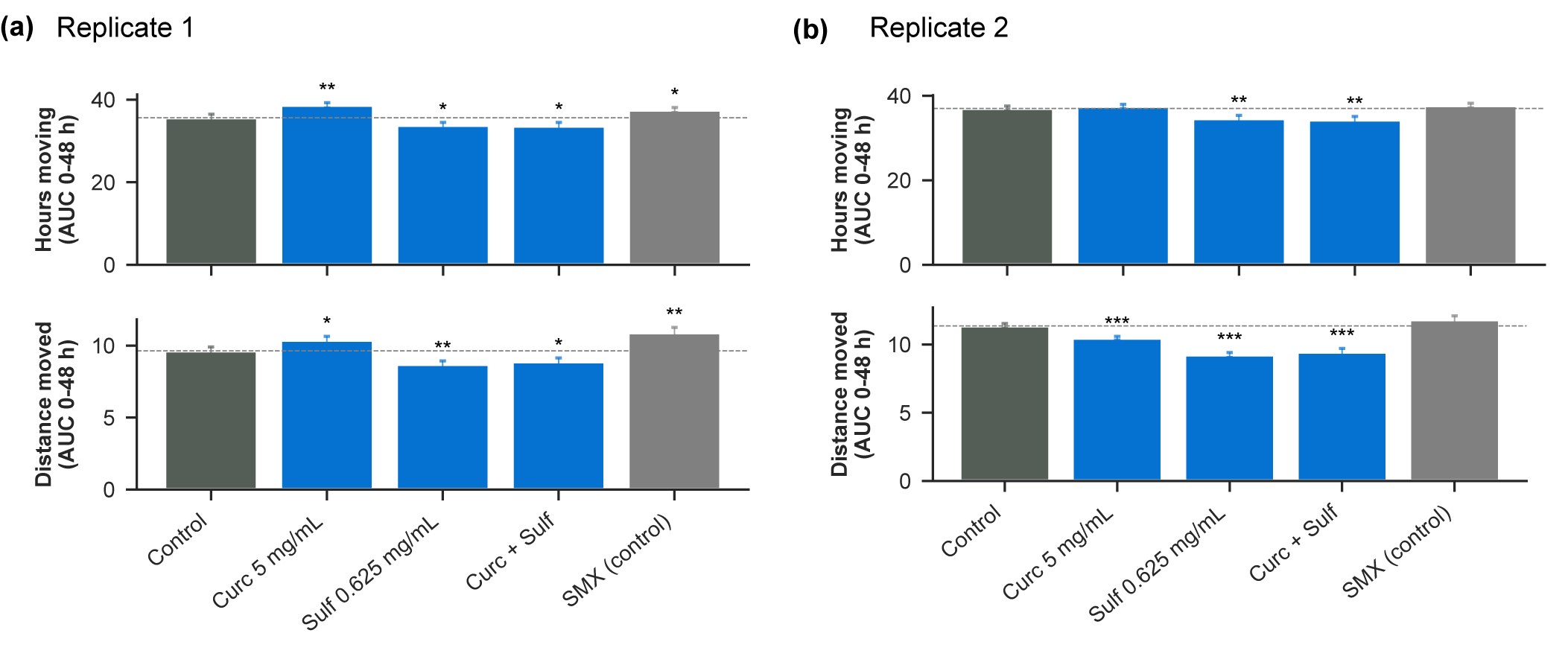

**Figure S8**: Early-window AUC (Day 0–2). (a) Replicate 1 and (b) Replicate 2. Bars show hours moved (top; AUC 0–48 h) and distance moved (bottom; AUC 0–48 h) for Control, Curcumin 5 mg/mL, Sulforaphane 0.625 mg/mL, Combination, and SMX (platform control). Values are mean ± SEM (technical). Stars denote a one-sided test on ΔAUC (treatment vs matched control): * p < 0.05, ** p < 0.01, *** p < 0.002; NS otherwise. Primary inference in the main text uses AUC Days 2–7 (see Figure 2 and Figure S7).

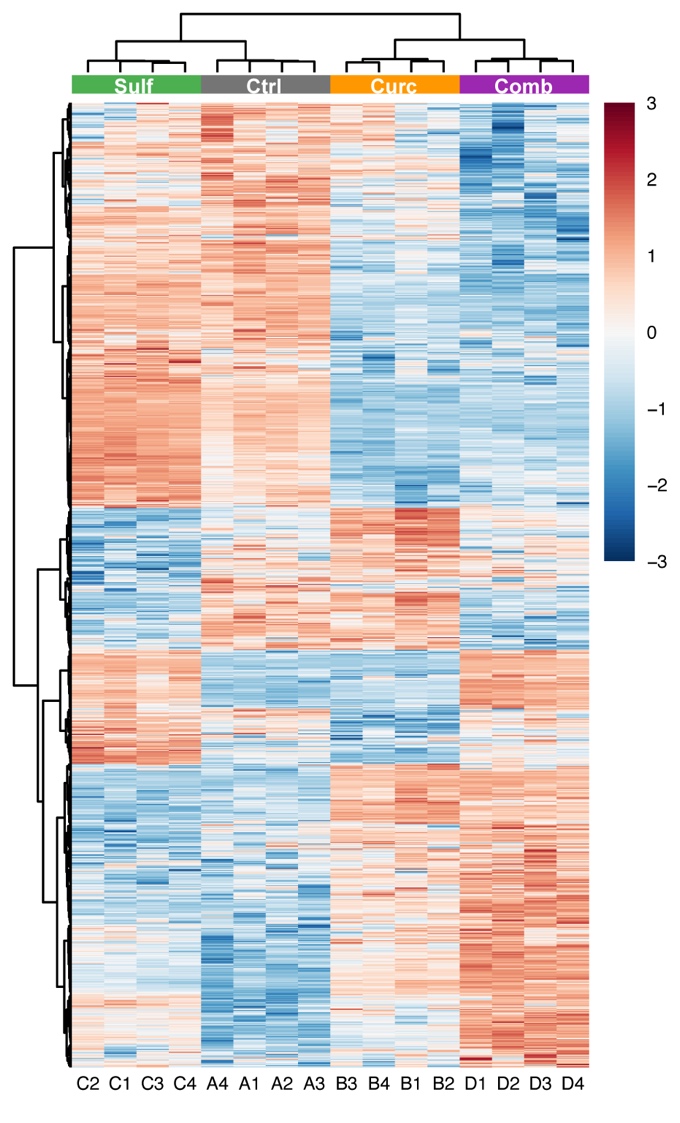

**Figure S9:** Heatmap of transcriptome-wide expression. Columns are the 16 libraries grouped by treatment (Sulf, Ctrl, Curc, Comb; colored chips above). Values are row-wise z-scores of log2-FPKM. Rows and columns were clustered using Euclidean distance with ward.D2 linkage. The heatmap shows distinct global profiles for each treatment and coherent clustering of biological replicates.

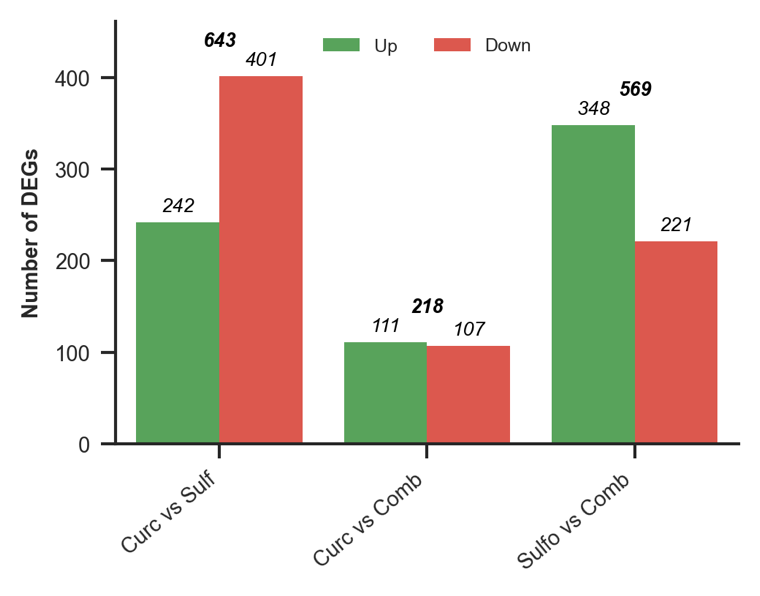

**Figure S10**: Pairwise DEGs between treatments (*padj* ≤ 0.05; |log₂FC| ≥ 1). Bars show numbers of Up and Down DEGs for Curc–Sulf, Curc–Combo, and Sulf–Combo; totals are annotated above each pair.

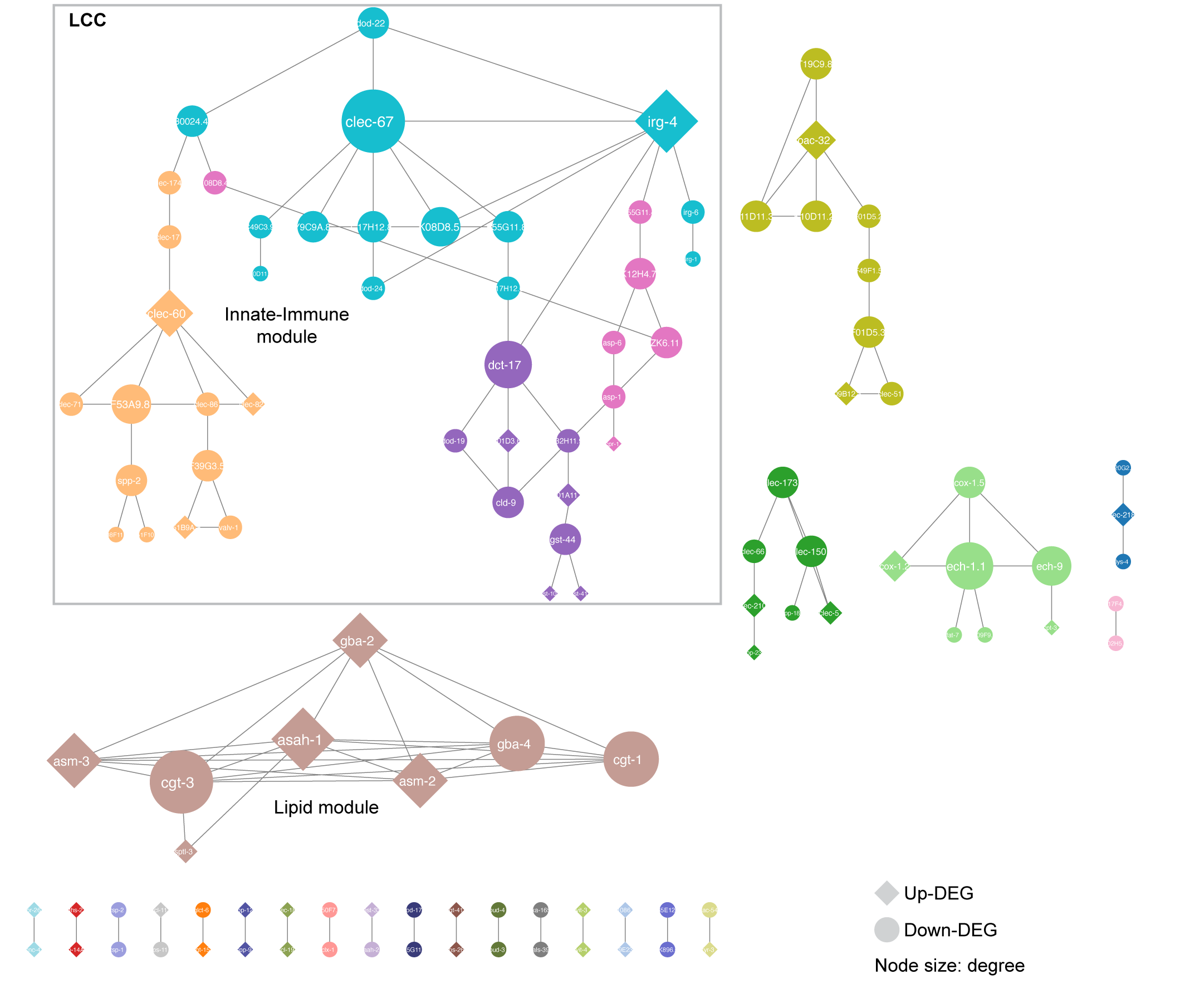

**Figure S11**: Complete STRING protein-protein network of DEGs under Curcumin treatment. Edges: STRING v12.0, combined score ≥ 0.7; nodes are DEGs only (*padj* ≤ 0.05; |log2FC| ≥ 1); isolates removed; communities by Glay (node color = community; diamond node = Up-DEG; circle node = Down-DEG, node size = degree of the node).

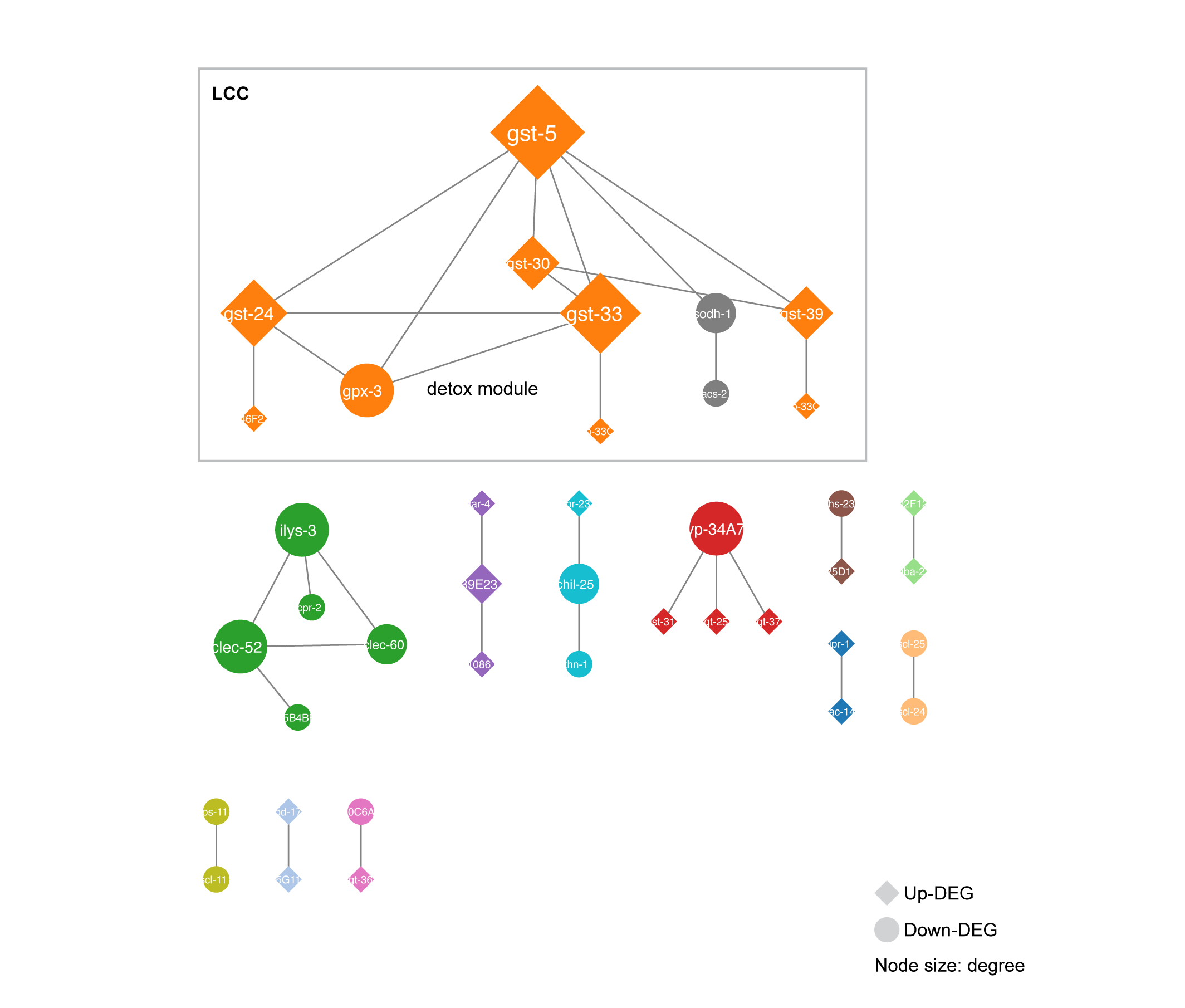

**Figure S12**: Complete STRING protein-protein network of DEGs under Sulforaphane treatment. Edges: STRING v12.0, combined score ≥ 0.7; nodes are DEGs only (*padj* ≤ 0.05; |log2FC| ≥ 1); isolates removed; communities by Glay (node color = community; diamond node = Up-DEG; circle node = Down-DEG, node size = degree of the node).

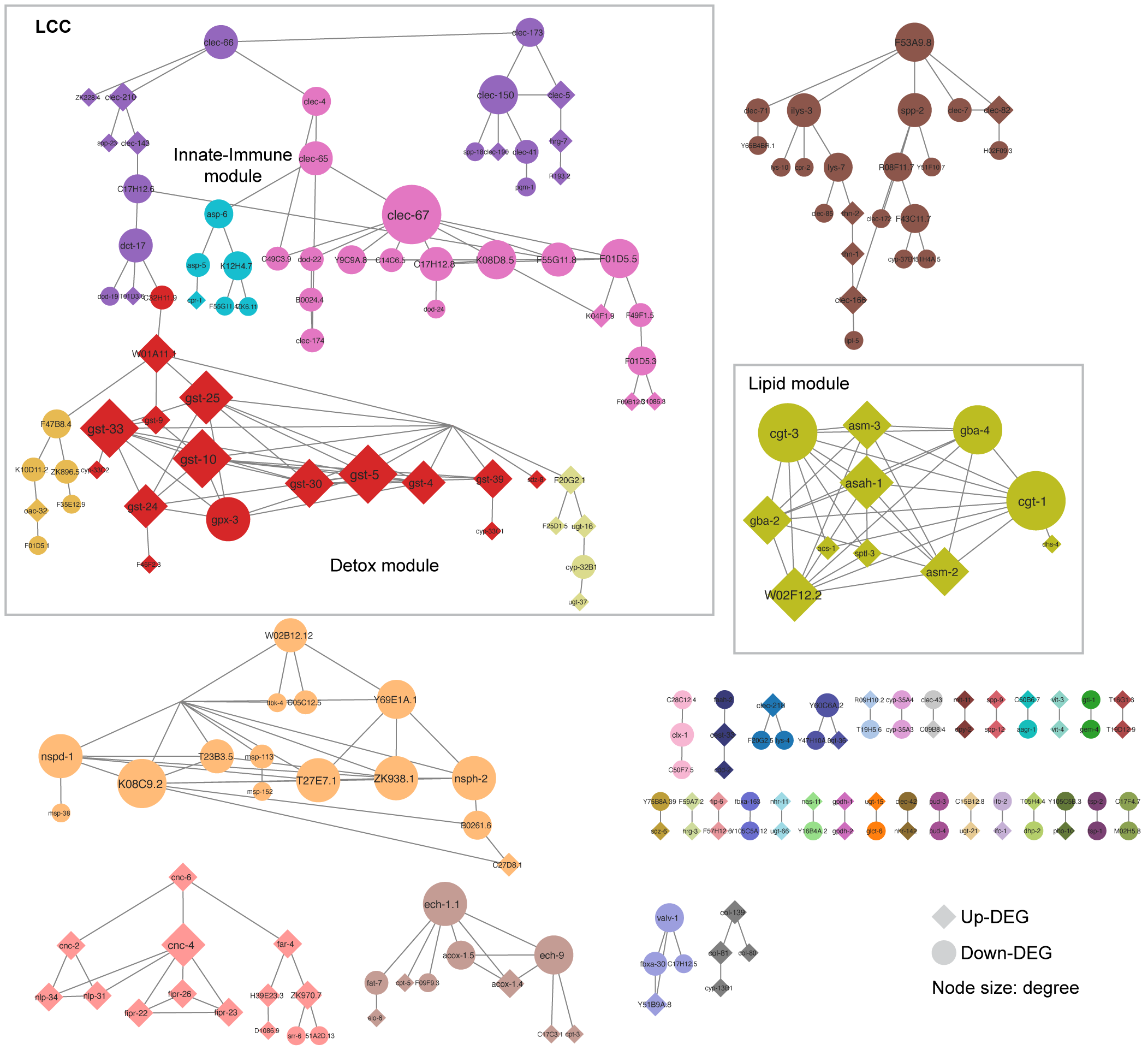

**Figure S13**: Complete STRING protein-protein network of DEGs under Combination treatment. Edges: STRING v12.0, combined score ≥ 0.7; nodes are DEGs only (*padj* ≤ 0.05; |log2FC| ≥ 1); isolates removed; communities by Glay (node color = community; diamond node = Up-DEG; circle node = Down-DEG, node size = degree of the node).
